# Urban Cohabscapes: exploring European Co-Habitative landScapes diversity, in the ECOLOPES framework

**DOI:** 10.1101/2024.11.11.622937

**Authors:** Gabriele Oneto, Katia Perini, Maria Canepa, Victoria Culshaw, Wolfgang Weisser, Anne Mimet

## Abstract

Urban liveability is closely related to the landscape features that support human societies. The promotion of liveability influences the well-being of both human and non-human inhabitants of the city, as the two share the same urban habitat. In this paper, we propose a classification of urban landscapes tailored to facilitate the acquisition of interdisciplinary knowledge on the multi-inhabitant liveability of urban landscapes. Our classification represents the layered urban dimensions through four different subclassifications: local and landscape Urban Form, Anthropic Imprint, and Biophysical Conditions. We developed a flexible and scalable methodology that enables us to expand to other cities, by using only opensource geospatial and remote sensing data. We modelled our four subclassifications at 10-metres through a set of automatic pipelines, running Principal Component Analysis as dimensionality reduction and KMeans unsupervised classification. In this paper we explore the application of our methodology in the framework of ECOLOPES, a project that investigates the human and non-human liveability in cities through the computational design of green building envelopes. We apply the methodology to three distinct European cities, i.e. Vienna, Munich, and Genoa. We produced a set of 12 raster maps from 64 variables, with a total of 32 classes and 1264 possible combinations. We analysed inter and intra-urban class frequencies by highlighting the primary spatial signatures over each sub-classification, and between the overall Functional Urban Areas and the core areas. In this paper we discuss how this approach could foster multidisciplinary studies on urban liveability that hold into account not just humans, but all living inhabitants in cities. From this application, we envision a second step where our methodology will be applied to the full extent of European cities.

## 1. Introduction

Human well-being and health are closely linked to the natural environmental systems on which human societies depend (Betley et al., 2011; Betley 2023). The One-Health perspective, which views humans as part of the planetary system, replaces the post-industrial connotation of humans as separate from nature (Felappi et al., 2020). This perspective aligns with the requirements of sustainable development encompassing all organisms, safeguarding life on Earth amid global changes (Audusseau et al., 2024). European cities are gradually adopting this view, moving away from designs and decisions that kept nature distant (Shingne, 2020). One example is Nature-Based Solutions (NBS, Eggermont et al. 2015), through which cities have developed innovative ways to enhance urban resilience and urban liveability for humans and biodiversity encompassing also animals by inserting natural elements, in particular plants into the urban fabric (Sowińska-Świerkosz and García, 2022). NBS are responsible for significant progress in terms of climate adaptation and biodiversity in cities, but researchers are signalling a limitation in comparing the NBS benefits for measuring human and non-human liveability (Apfelbeck et al., 2020). Overcoming this limitation would require, at the fundamental knowledge level, a comparative tool capable of linking urban landscapes to the different types of inhabitants it supports (Hunter et al., 2023, Potter et al., 2023). Such a comparative tool would help coordinate the different professionals involved in designing and planning NBS, such as architects, urban ecologists, planners, and social scientists (Broman and Robèrt, 2017; Khan et al., 2020). In this paper, we consider using multiple interconnected classifications of urban landscapes for addressing this comparison limitation.

The urban landscapes features that influence the coexistence of human and non-human inhabitants are both biotic and abiotic factors (Aronson et al., 2016; Mitchell and Devisscher, 2022). Primarily, regional climatic and biogeographical factors determine the proliferation of particular species, and their capacity to foster in a set urban landscape (Aronson et al., 2016). Coexistence, or co-habitation since the different species share the same urban habitation space, is also influenced by other factors. Different urban landscapes have each grown into supporting their inhabitants by defining their own human-centric services, by adapting the urban form, and from being influenced by socioeconomic and cultural factors (Alberti et al., 2020; Bettencourt, 2021). Continuing, urban landscapes support species interactions through an extensive list of shared processes, such as the need for resource gathering (e.g., nutrient cycling, healthcare), adapting to abiotic features (e.g., air purity, shelter and housing), and moving through the landscape for sustaining themselves (e.g., pollination, work-commuting) (Alberti, 2023). The diversity of urban landscape features, each corresponding to a different urban dimension, can be explained by status of layered information that each urban landscape has undergone to further its liveability (Aronson et al., 2016). This layered vision can be used to define the liveability dimensions of urban landscapes for the different inhabitants, or stakeholders, interactions, through the lens of abiotic features, physical built and natural features, and cultural and social-induced features (Baccini and Brunner, 2012; Egerer et al., 2021, McPhearson et al., 2022). This approach streamlines the process of analysing stakeholder’s interactions by obtaining urban landscape conditions from its features (Hölscher and Frantzeskaki, 2021; Verrelli et al., 2022). We use this layered structure to build our classification, and we refer to existing studies to define the requirements to effectively represent the stakeholder interactions in term of extension and flexibility of application, resolution of usage, and generalisation through different urban conditions (Shingne and Reese, 2022).

Previous studies (Li et al. 2019; Liu et al., 2021; Paumelle et al., 2023) have shown that classifications can effectively describe urban landscape conditions through landscapes features, while catering for both human and non-human inhabitants. Typically, these classifications use opensource databases (for example, Copernicus High Resolution Layers, and Global Human Settlement Layer) and traditional clustering algorithms (for example, hierarchical clustering, and KMeans), meaning that these classifications are cost-effective opportunities and statistical methodologies for characterizing entire urban areas on a large extent. Numerous classifications or frameworks exists to describe urban areas from landscapes features (Zhang et al., 2023), most focusing on connecting climatic and biophysical features to urban morphology (Joshi et al., 2022; Lentini et al., 2024). These classifications exhibit a crescent trajectory of development across different levels of structural complexity, ranging in term of finer resolution, larger flexibility of application, and increased generalisation. Simply put, the main factor is the translation of the urban dimensions into how the classification is structured, resulting in a different desired level of detail (Bollinger et al., 2005). Regarding generalisation, classification can choose between adhering to a species-specific point of view, for example by adopting variables that describe species occurrences, or a generic point of view, that opt to select information valid for all stakeholders (Bollinger et al., 2005). Generic classifications are valued for their flexibility and heterogeneous description of the urban dimensions, and are used in climate-change assessment (Mokhtari et al., 2022); urbanization and land cover analysis (Mendoza Beltran et al., 2023), land cover change assessment (Luo et al., 2020), biodiversity analysis (Li et al., 2019), green infrastructure planning (Morpurgo et al., 2023), and urban landscape characterization (Stokes and Seto, 2019). These classifications make explicit the urban heterogeneity by accounting for different urban dimensions, such as physical features and socioeconomics, without sacrificing resolution and scale. However, these generic classifications predominantly adopt a human-centric perspective when describing the urban environment, and often overlook another kind of generalisation, i.e. a non-species-specific point of view that opt to select information valid for all stakeholders. Species-generality, or neutrality, was introduced to remove not only the human-centric perspective bias prevalent in generic classification, but also species-specific variables, such as density or connectivity of a particular animal or plant (Mimet et al., 2013, 2014; Marrec et al., 2020). Species-generic urban classifications have been successfully used in studies such as multi-risk assessment analysis (Beevers et al., 2022), smart city applications (Majchrowska et al., 2022), and environmental and agricultural analyses (Halstead et al., 2021). Creating an urban landscape classification tailored to serve interdisciplinary research on urban liveability requires to combine the best features of the species-generic urban classifications, with enough spatial extent, flexibility of application and resolution to compare different conditions.

In this paper, we propose a classification of the urban landscapes tailored to describe the diversity of urban landscapes, facilitating the acquisition of interdisciplinary knowledge on the liveability of human and non-human inhabitants. In this perspective, the proposed classification meets the following criteria:

- Represent the stratification of the urban dimensions, using different harmonised layers and heterogeneous variables;
- Use modelling criteria and variables that account for the liveability of different types of inhabitants, or stakeholders, referring to existing variables found in the literature;
- Develop a flexible and scalable methodology ensuring that the classification could be expanded and applied to any European city to enable between-cities comparisons, by using opensource geospatial and remote sensing data and selecting variables that could be feasibly obtained in different regions;
- Model data at high resolution (10-metres) to capture urban landscape heterogeneity and enable within city comparisons, diversifying for the difference in scopes for each variables by using a multiscale approach; We then apply our classification to three distinct case studies European cities: Vienna (Austria), Munich (Germany), and Genoa (Italy).

The proposed Cohabscape (Co-Habitative landScapes) methodology delineates urban landscapes for multiple inhabitants through four distinct sub-classifications representing urban dimensions, each with a similar computation. The first and second sub-classification, **Urban Form**, defines at the local scale physical features that shape urban morphology and its diversity; and at the landscape scale by analysing spatial configuration patterns (Krueger et al., 2022). The third sub-classification, **Anthropic Imprint**, deals with collecting human leftover interactions within urban areas, both actively and passively, such as through land use policies and anthropic emissions, including services and economic orientation (Schwarz, 2010). The fourth sub-classification, **Biophysical Conditions**, refers to abiotic attributes that characterise cities independent of internal city dynamics. This last sub-classification covers environmental, hydrologic, orographic, and climatic top-down conditions that influence urban landscapes.

## 2. Materials and methods

### 2.1 Overall Methodology

The Cohabscape methodology consists of four steps, i.e. defining the urban boundary, forming the dataset for each sub-classification, preprocessing, and unsupervised clustering. The first step regards setting the computation spatial limits (i.e., the urban boundary) and justification for the resolution (Fig. 1.1). The second step address selecting relevant variables for each sub-classification, the selection of opensource data repositories and computation, between locally-representative and at the landscape scale (Fig. 1.2). The third step preprocess data in terms of standardization techniques and dimensionality reduction to simplify and reduce the number of variables to smooth the classification (Fig. 1.3). The fourth and last step explains the clustering approach through unsupervised classification, optimisation and cluster content analysis (Fig. 1.4). Finally, we describe our case studies and the information we will extract from our classification (Fig 1.5).

**Fig. 1.**
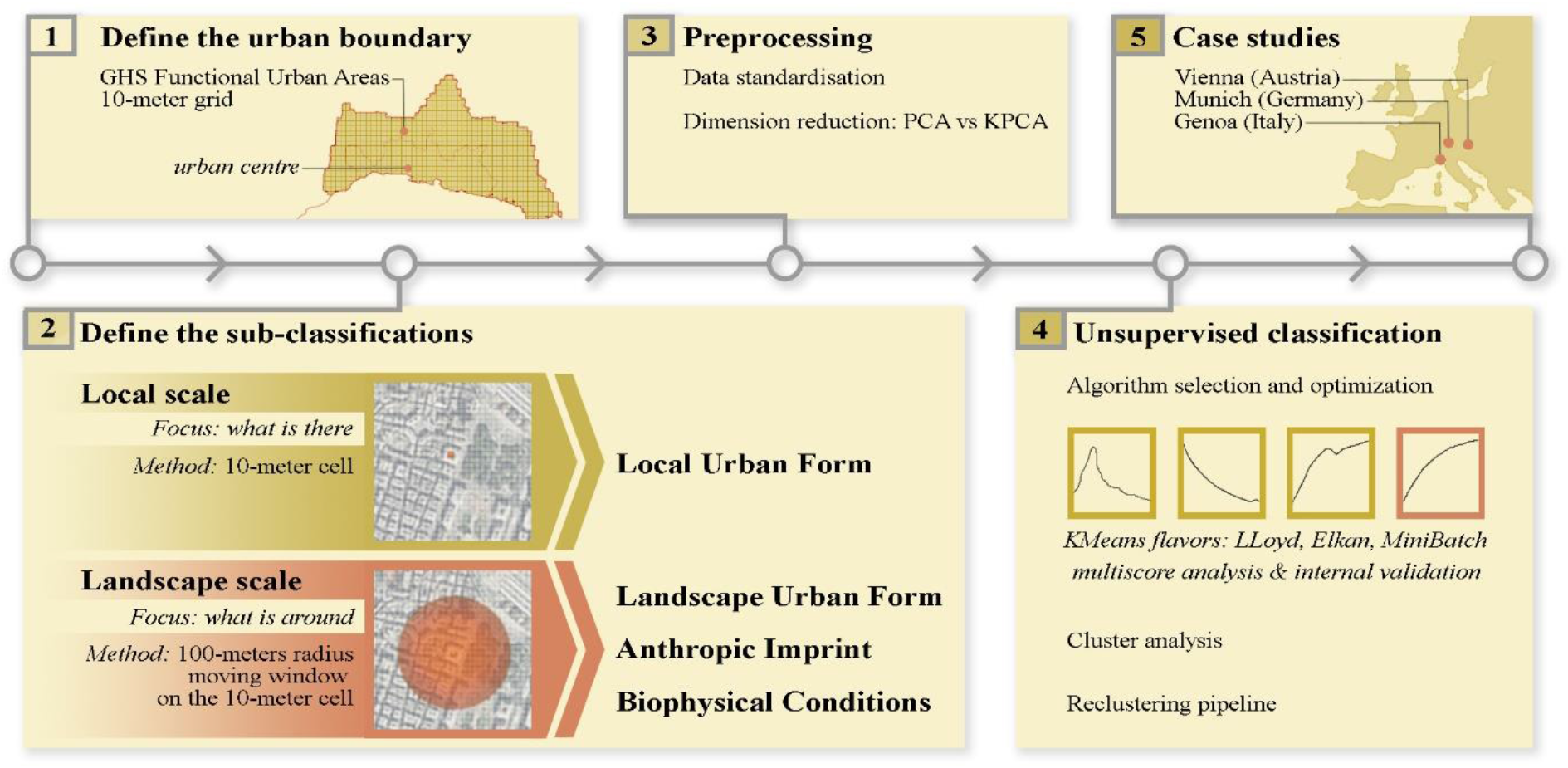
1 to 1.5: Steps of the Cohabscape methodology. Step 1 defines the urban boundary and global settings of the framework. Step 2 defines the sub-classification pipelines, their variables and the different scope: local vs landscape. Step 3 refers to the preprocessing and dimensionality reduction. Step 4 is the classification and score analysis. Step 5 refers to the application to the three case studies.

Annex A covers in full the variables and methodologies used to compute the dataset for each sub-classification. Annex B explain the preprocessing through data standardization and dimensionality reduction techniques. Annex C contains technical information on the unsupervised clustering and internal validation. Annex D combines results from Annex B and C to name the classes through a data driven methodology.

### 2.2 Step 1: Defining the urban boundary

Within this study we limited the computation extent on urban areas by adhering to the Global Human Settlement Layer Functional Urban Areas (Schiavina et al., 2023), promoted by the Organization for Economic Co-operation and Development (https://www.oecd.org/en.html) and the European Union (Dijkstra et al. 2019). We use an equiareal 10 × 10 metre global grid projected onto the Mollweide (ESRI:54009) coordinate system (further details are in Annex A).

### 2.3 Step 2: Defining the sub-classifications

The variables were chosen to reflect the aforementioned urban dimensions: urban form (i.e., physical features and landscape patterns), anthropic imprint (i.e. leftover human interactions), and biophysical conditions (i.e. climate and geographical features). We considered in our selection also variables that are known to influence or be influenced by other variables, as their inherent collinearities would have been addressed in a later stage. Each variable featured uniquely in a different sub-classification, without repetitions. We approached the selection of variables with two criteria. Firstly, we wanted to maximise global applicability for our methodology, meaning selecting variables that could be feasibility obtained (or calculated) in different world regions. Secondly, we based our selection on the literature, i.e. variables used in other classifications and categories of variables that are identified in the literature to be important for the human and non-human inhabitants.

In the case of Urban Form at local scale, both built, and vegetation conditions were considered, which gave a total of nine variables (Table 1). At the local scale, urban form can be described by variables such as artificial features that predominantly impact landscape definitions for built area, volume, and infrastructure (Seto and Fragkias, 2005, Boeing, 2018). Similarly, urban form can be described by variables of natural features (Wentz et al., 2018).

**Table 1.**
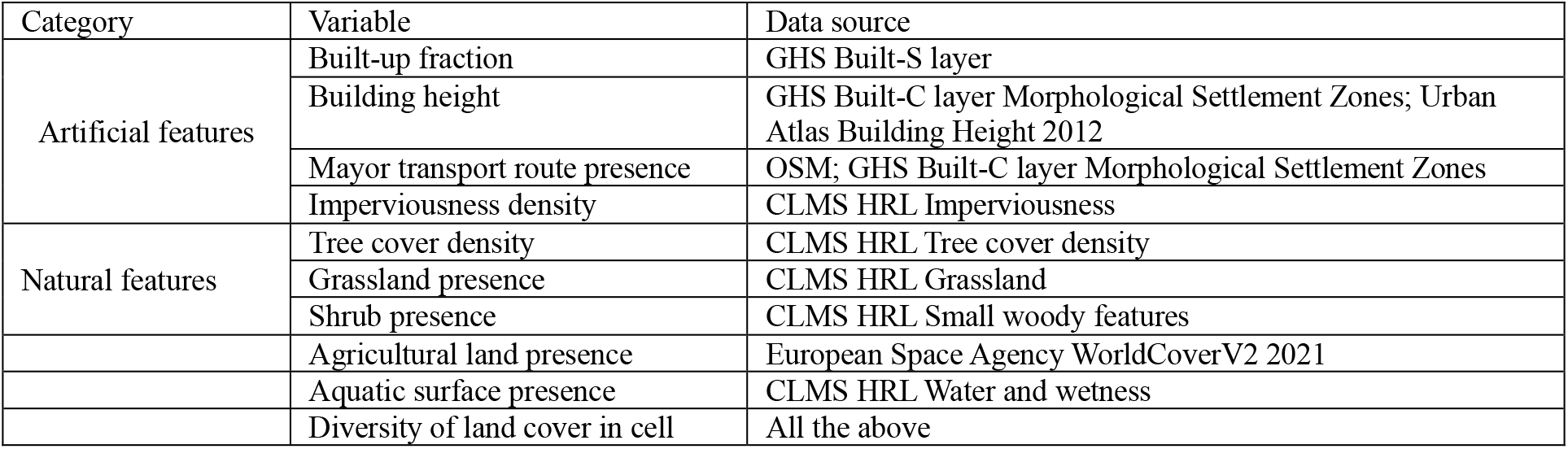
Variables for Urban Form local sub-classification. We use data that describes artificial (Global Human Settlement – GHS, Pesaresi et al. 2023; European Environment Agency – EEA, 2022; Open Street Maps – OSM, www.openstreetmap.org) natural features and diversity (Copernicus Land Monitoring Service High Resolution Layer – CLMS HRL, EEA, 2020a; EEA 2020b; EEA 2020c; EEA 2020d; EEA 2023; https://worldcover2021.esa.int; Zanaga et al., 2022). More details are in the additional material (Annex A).

The Urban Form at the landscape scale sub-classification builds upon the variables used in the local scale Urban Form sub-classification and is based on 24 variables. In this second Urban Form, we considered both composition (for each class) and configuration (for all classes combined) variables. The composition variables were obtained by firstly resampling at 10-metres, then calculating the mean and standard deviation of each principal component axis present at the local scale, within 100-metres moving window, for a total of 22 variables. Through configuration variables, we studied landscape patterns by measureing aggregation with contagion, and diversity via Shannon’s Diversity Index, using the same radius of moving window used for the composition variables. We used the stand-alone software FragStats 4.2 (https://fragstats.org/; McGarigal K. et al., 2023). Notably, Urban Form at landscape scale share variables with the local scale but is distinguished by scale and computational pipeline to make sure the final information is different while remaining ecologically and urbanistically valid (Clifton et al., 2008; Schwarz, 2010; Fleischmann et al., 2021).

For the Anthropic Imprint sub-classification, we considered demographic, land use and anthropic emissions information for a total of 13 variables (Table 2). All listed variables were resampled at 10-metres and computed within a 100-metres moving window at 10 metres resolution. Demography and land use directly represent human activity in urban areas. Whereas the former is a more direct representation, the latter describes passive human activities in terms of capacity of operating in urban landscapes, such as very intensive industrial operations or park regulations (Clifton et al., 2008). Additionally, passive anthropic emission, such as night time light pollution and heat emissions, are known to produce effects in urban landscapes (Crippa et al., 2021).

**Table 2.**
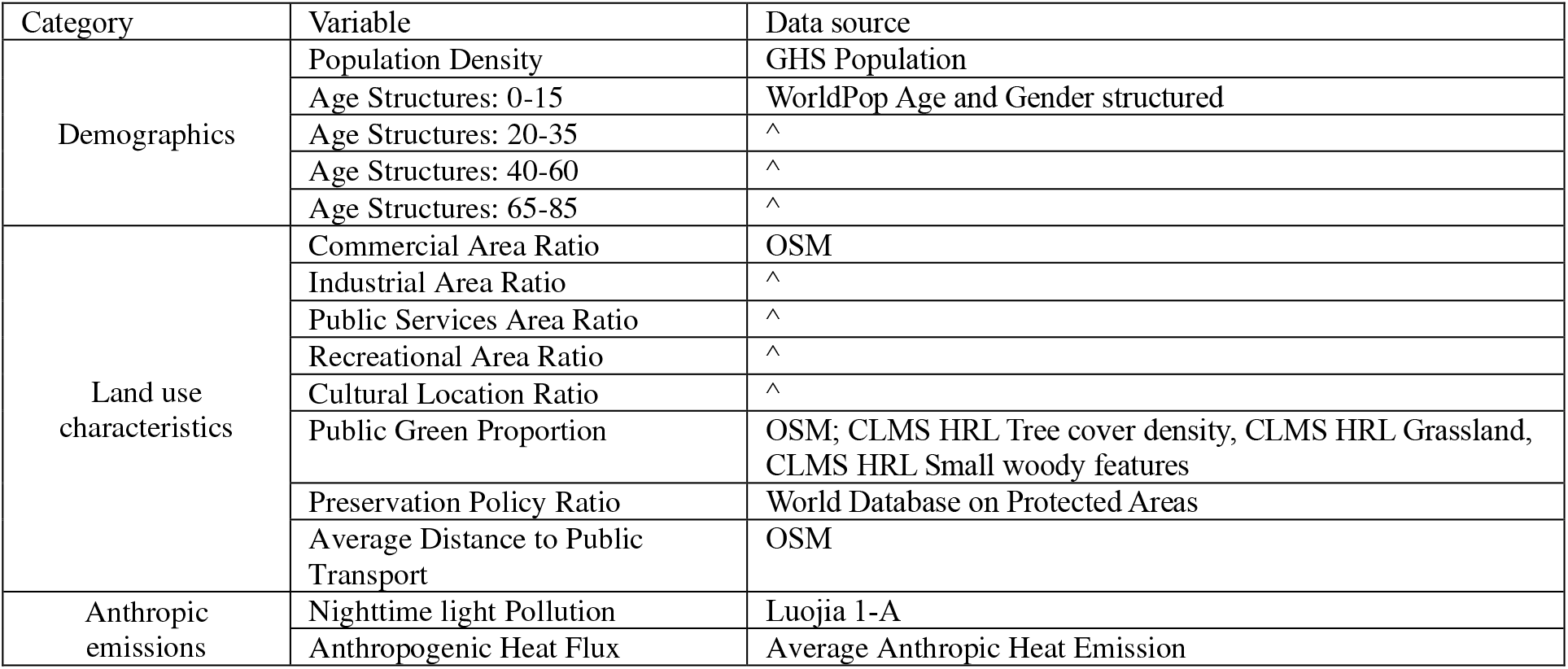
Variables for Anthropic Imprint sub-classification. We use data that describes demography (Global Human Settlement – GHS, Schiavina et al., 2023; Bondarenko et al. 2020), land use characteristics (Open Street Maps – OSM, www.openstreetmap.org; Copernicus Land Monitoring Service High Resolution Layer – CLMS HRL, European Environmental Agency - EEA 2020b; EEA 2020c; EEA 2020d; https://www.protectedplanet.net/en; United Nations Environment Programme - World Conservation Monitoring Centre, 2019), and anthropic emissions (Chen et al., 2022; Varquez et al., 2021). More details are in the additional material (Annex A).

Finally, the Biophysical Conditions sub-classification was composed by 19 variables divided between hydrology, topography, soil characteristics, and climate (Table 3). For this fourth sub-classification, originally, we considered using climate classifications, such as ones based on the Köppen-Geiger criterion (Beck et al., 2018), however we noted a lack of definition provided by this criterion within the city. We decided, instead, to represent regional climatic and geographic features through individual abiotic indicators by extracting both absolute and relative climate variables, with the latter being divided by the mean values for each city. We disregarded standard deviation for climate data because our preliminary results demonstrated the coarse native resolution of 1000-metres meant a strong gridded pattern that negatively influenced the clustering. Additionally, we computed the adjusted mean values for each climate variable, by dividing each by the local mean value in order to homogenise the comparison between potentially vastly different geography regions. All data was resampled at 10-metres and computed through a 100-metres moving window.

**Table 3.**
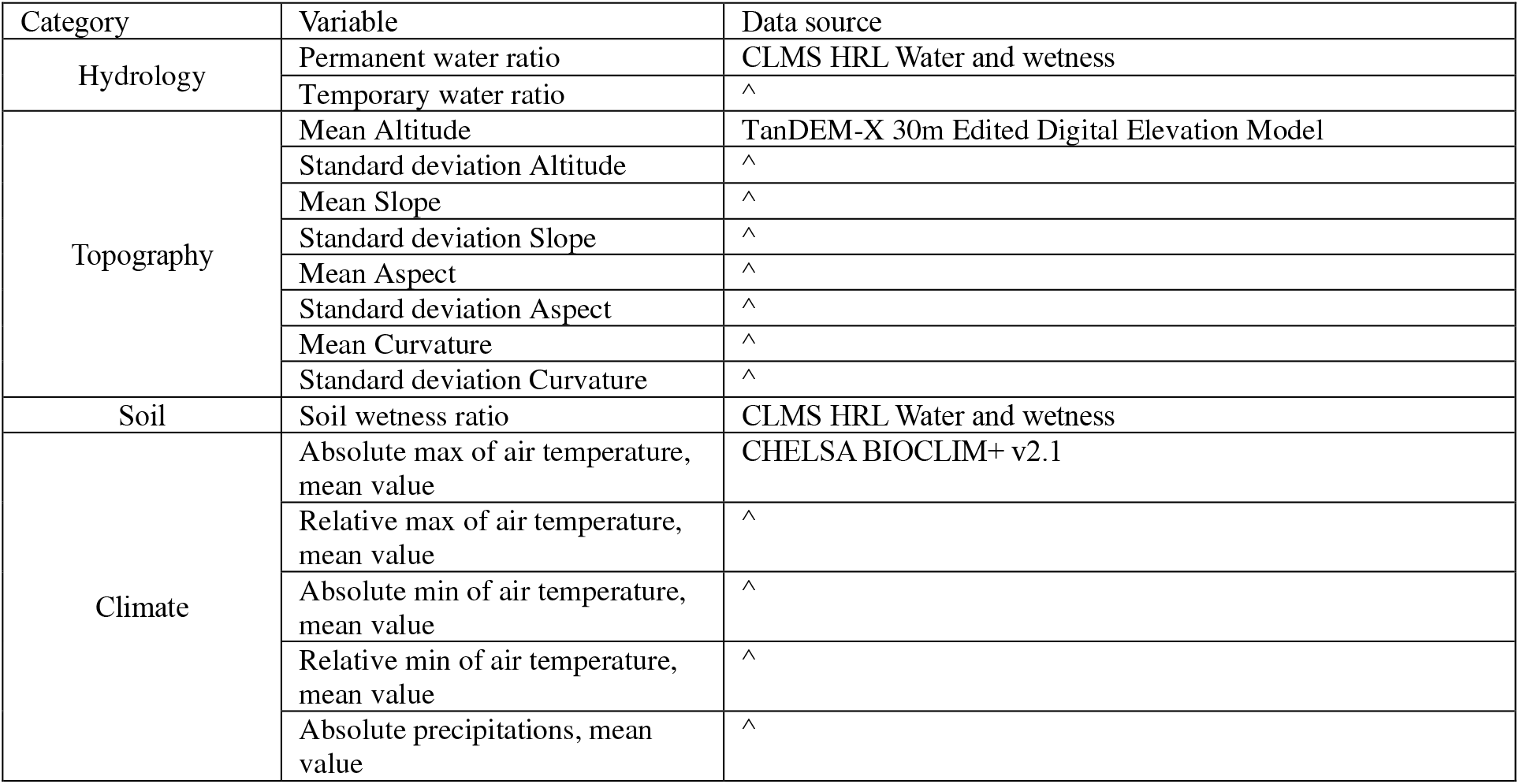
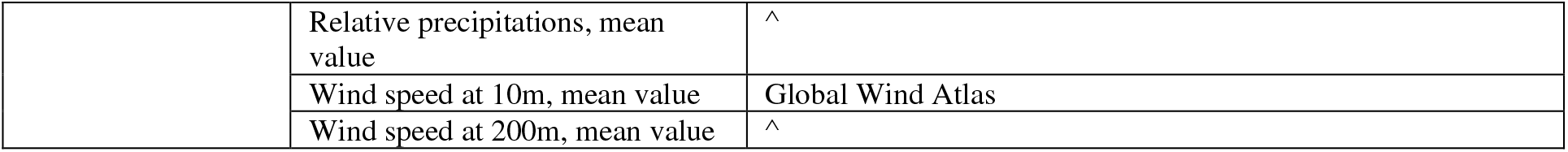
Variables for Biophysical Conditions sub-classification. We use data that describes hydrology (CLMS HRL, European Environmental Agency - EEA 2020d), topography (González et al., 2020), soil wetness (CLMS HRL, European Environmental Agency - EEA 2020d), and climate (Karger et al., 2017; Davis et al., 2023). More details are in the additional material (Annex A).

| Category | Variable | Data source |
| --- | --- | --- |
| Hydrology | Permanent water ratio | CLMS HRL Water and wetness |
|  | Temporary water ratio | ^ |
| Topography | Mean Altitude | TanDEM-X 30m Edited Digital Elevation Model |
|  | Standard deviation Altitude | ^ |
|  | Mean Slope | ^ |
|  | Standard deviation Slope | ^ |
|  | Mean Aspect | ^ |
|  | Standard deviation Aspect | ^ |
|  | Mean Curvature | ^ |
|  | Standard deviation Curvature | ^ |
| Soil | Soil wetness ratio | CLMS HRL Water and wetness |
| Climate | Absolute max of air temperature, mean value | CHELSA BIOCLIM+ v2.1 |
|  | Relative max of air temperature, mean value | ^ |
|  | Absolute min of air temperature, mean value | ^ |
|  | Relative min of air temperature, mean value | ^ |
|  | Absolute precipitations, mean value | ^ |
|  | Relative precipitations, mean value | ^ |
|  | Wind speed at 10m, mean value | Global Wind Atlas |
|  | Wind speed at 200m, mean value | ^ |

Data was collected both manually, automatically and through distributed API where available (OSMNX, Boeing 2024; chelsa_cmip6 package; Karger et al., 2017). Data was then reprojected onto our Mollweide grid and computed at different scales. The local scale used for Urban Form layer operated cell-by-cell. The landscape scale, used for the remaining three sub-classification, used a rounded 21 cell-diameter window centred on each cell. Additional information regarding the individual methodologies and pipelines can be found in the additional material (Annex A).

### 2.4 Step 3: Preprocessing and dimensionality reduction

For each sub-classification, we merged together the heterogeneous data masking off pixels outside the buffered Functional Urban Areas. We obtained pixel coordinates by loading each city and saving the coordinates separately, referring to a ubiquitous index. Since our data would then be later sampled and pre-processed, we could reconstruct the original position of each cell and reform the data stacks.

For the pair of urban form layers, we deviated slightly and extract water pixels from the starting batch, preferring to represent water unambiguously. We removed all pixels where water was present, preserving a stable layer that would be then added to the local classification at the end. By doing the same process at a later stage, we produced water patches in the landscape scale similarly.

We then proceed to reduce collinearity and simplify the variance by using feature reduction techniques, that are shown to improve clustering performance (Li et al., 2019; Paumelle et al., 2023). We tested two alternative approaches for dimensional reduction. On one hand, we run Principal Component Analysis (PCA), estimating the number of components by looking at the minimum number of eigenvalues needed to explain 95% of the total variance. On the second hand, we also explored the performance of Kernel Principal Component Analysis (KPCA). We run through a grid search of the best kernel and hyperparameters, choosing between linear, polynomial, radial basic function, and additive chi squared, and modelling 50 components with the Nistroem sklearn component. Both analyses were conducted using the sklearn package with the PCA method, nystroem kernel approximation, MinMaxScaler, and gridsearchcv pipelines.

After each analysis was completed, we plotted the results and confronted the normal PCA with 0.95 explained variance with the best KPCA approximation results, and selected the one that could describe the dataset by using less axis to explain the same variance.

### 2.5 Step 4: Unsupervised Classification

#### 2.5.1 Algorithm selection and optimisation

After the dimensionality reduction, we studied the appropriate clustering algorithm and best number of classes. After preliminary tests, we landed on using a set of KMeans flavors of algorithm, limiting the optimisation between the Lloyd, Elkan and MiniBatch, using sklearn implementations for all three. We favoured the ease of computation thank to the fact that the spatial extent and resolution resolved in over 60’000’000 cells, albeit the sub-classification datasets presented few and collinear variables. The alternatives are considered in the additional material (Annex C).

The evaluation of the performance of the algorithms was done with the Silhouette scores as the primary indicator for the best internal validation and added the Calinski-Harabasz and Davies-Bouldin as secondary indicators (Palacio-Niño and Berzal, 2019). We additionally included the plotting of inertia to compare the results with the Elbow method as an internal validation criterion.

We run the different flavours of KMeans through a range of 6-30 possible number of classes (k), plotting the different scores. Our selection was based on the comparative performances of the three different scores, keeping the Elbow Method as an analogic indicator of good estimation. We used a three-fold multicriteria method for comparison. We compared each combination of scores by looking at the difference between the scores value at any k and the scores value of the best solution. By summing the differences for each k value, we obtain the k value that would come closer to most of the best scores.

#### 2.5.2 Cluster analysis

We explained the content of the classes by referring to the Principal Component values for each class (Fig. 2). Through a boxplot visualization, we plotted the normalized Principal Components from -1 to 1. We then divided the data spread for the box values and median in high (>|0.6|), medium (from |0.6| to |0.3|) and low (from |0.3| to |0.1|). This approach directly linked the Principal Component box and median values to a particular covariance to the original variables, indicating which class were strongly influenced by which variables. We especially tailored the analysis to explain classes through axis with narrow box distribution. A full analysis of the boxplots and reasoning is contained in the additional material (Annex D).

**Fig. 2:**
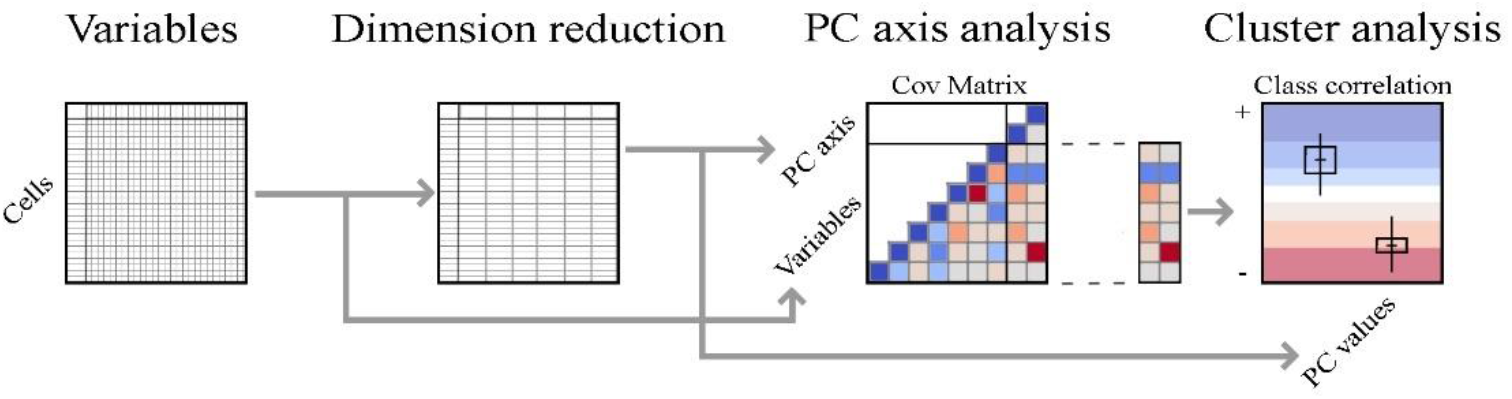
The cluster analysis represents each class content by referring to the correlation values between each principal component and the variables. After feature reductions, the principal components and variable covariance is calculated. These values, split between high, medium and low correlation, are visualised through boxplots for each class.

After naming the classes, the values were remapped to improve readability (Fig. 2).

#### 2.5.3 Reclustering pipeline

Given the broad size of our clustering area, i.e. the FUA, some inner-city variables could be synthetized in favour of larger scale peri-urban variables. To contrast this behaviour, we set up an additional pipeline to reprocess those classes we deemed could further explicit their content. This pipeline goes through a second round of unsupervised classification, by extracting specific variables from the original dataset. By selecting the variables that we want to better emphasize, we firstly identify the Principal Components with at least 0.5 absolute covariance with those variables. Then, for each class, for each selected Principal Components, we confronted the component’s median with the mean covariance between that component and its variables. If the median was closer to the mean covariance, the class was considered for the rerun. After selecting and extracting the pixels for all classes to rerun, the same algorithm optimization was run limiting the k range from the existing number of classes plus one and five times that amount. Finally, we merged the new and old classes. After preliminary results, we choose to deploy this pipeline only on the Urban Form local sub-classification, as the finer native resolution could produce additional insights.

### 2.6 Case study

The Cohabscape methodology was developed within the framework of the ECOLOPES project (https://ecolopes.org/), and was applied to the project European cases of ECOLOPES: Vienna (Austria), Munich (Germany), and Genoa (Italy) (Fig 3). The ECOLOPES project embodies a biophilic approach by offering a modelling framework for cohabitation strategies through architectural designs. This innovative approach seeks to revolutionize urban design, fostering inclusivity for multiple stakeholders including humans, plants, animals, and microbiota. At its core, an *ecolope* represents a building envelope - a computationally generated NBS – designed to sustain the lives of all these stakeholders, nurturing their flourishing and well-being as fundamental design objectives (Weisser et al., 2023). In the ECOLOPES initiative, while context-based ecological modelling and architectural parametrization cover the definition of the architectural shape, an open question remain on the landscape behaviour and the classification of the habitat in which an *ecolope* is located.

**Fig. 3:**
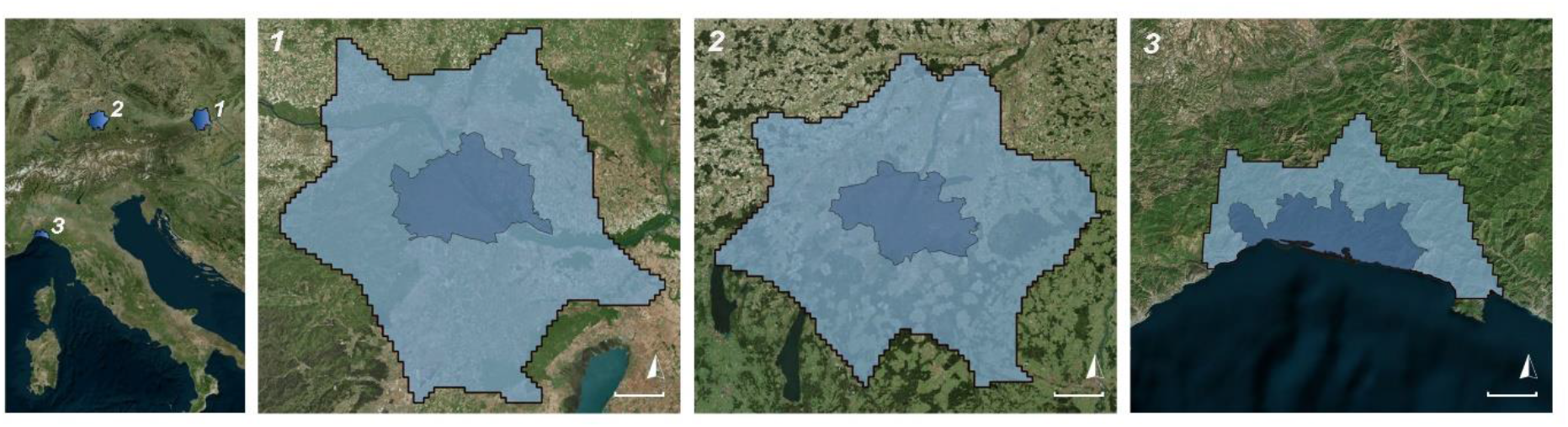
Localization of the three case studies (left to right 1 Vienna, 2 Munich, and 3 Genoa), highlighting the different analysed boundaries: the Functional Urban Area (FUA, in cyan), and the core city centre corresponding to the municipality (blue). The white scale bar corresponds to 10000-metres.

The three case studies allow to apply test the classification in different urban contexts, i.e. from different population densities, urban morphologies, geographical location, and climate (Fig. 4). Additionally, the case studies will be used to discuss what the classification could represent in term of interdisciplinary research and applications. Specifically, we focused on analysing the relationship between different sub-classification in the three case studies by obtaining quantitative measures on the frequency of occurrences of particular combinations of classes. We approached our analysis in two ways. Firstly, we confronted the sub-classifications separately city-by-city and understood the regionality for each class frequency. We used radar plots for describing the sub-classification distributions through the different cities. Secondly, we compared the cities together but looked at the distribution of classes frequencies between different areas of the FUA, i.e. the very urban centre corresponding to central municipalities and the remaining peri-urban areas. We defined the urban centre by using administrative municipality boundaries from OSM, and the peri-urban areas by the difference between the whole FUA and urban centre (Fig. 5). We plotted the results onto twelve distinct heatmaps, representing all possible pairwise combinations of sub-classifications between all cities.

**Fig. 4:**
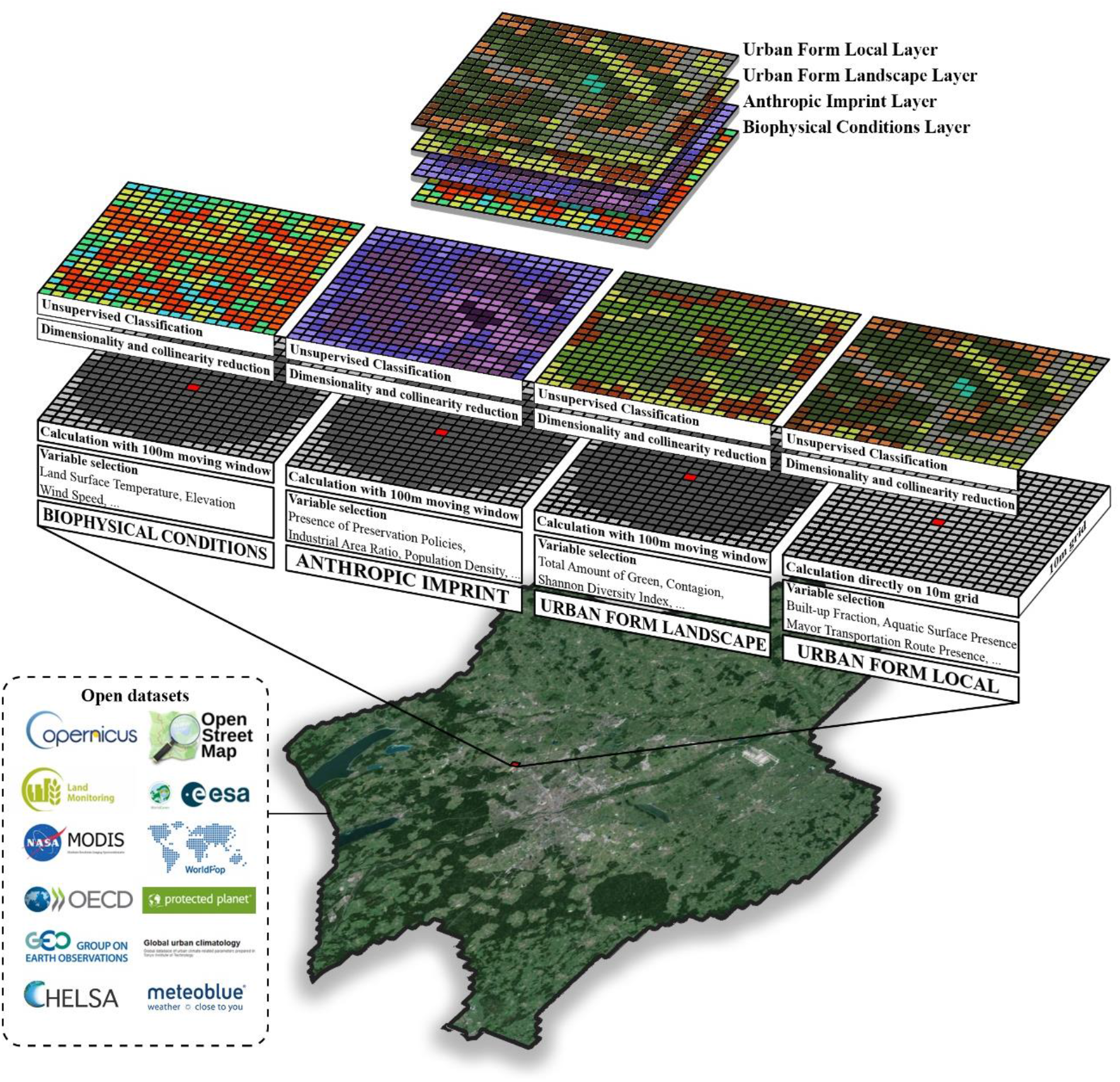
Application of the Urban Cohabscape classification to the Munich case study, from the bottom: definition of the city boundary through FUA, data collection and computation for each layer, dimensionality reduction and clustering, resulting in four rasters at 10 metres resolution.

## 3. Results

### 3.1 Obtaining the sub-classifications

From the variable selection, we retrieved opensource data and processed four different pipelines for all cities, each corresponding to an individual sub-classification. We integrated all cities data and run the preprocessing and classification for each sub-classification, the results of which are hereby presented.

***The Local Urban Form sub-classification*** was created by reducing the variables from nine down to six using KPCA. The unsupervised clustering was run two times, using the reclustering pipeline by extracting KPCA axis strongly correlated to artificial features. The two clustering were run using Elkan KMeans with k=10 and Elkan KMeans with k=4 by rerunning three classes, obtaining 13 distinct classes (Fig. 5.1). Among the classes, we noted a general prevalence of classes dominated by natural features (nine distinct classes) constrasting with fewer artificial dominated classes (four classes).

**Fig. 5.**
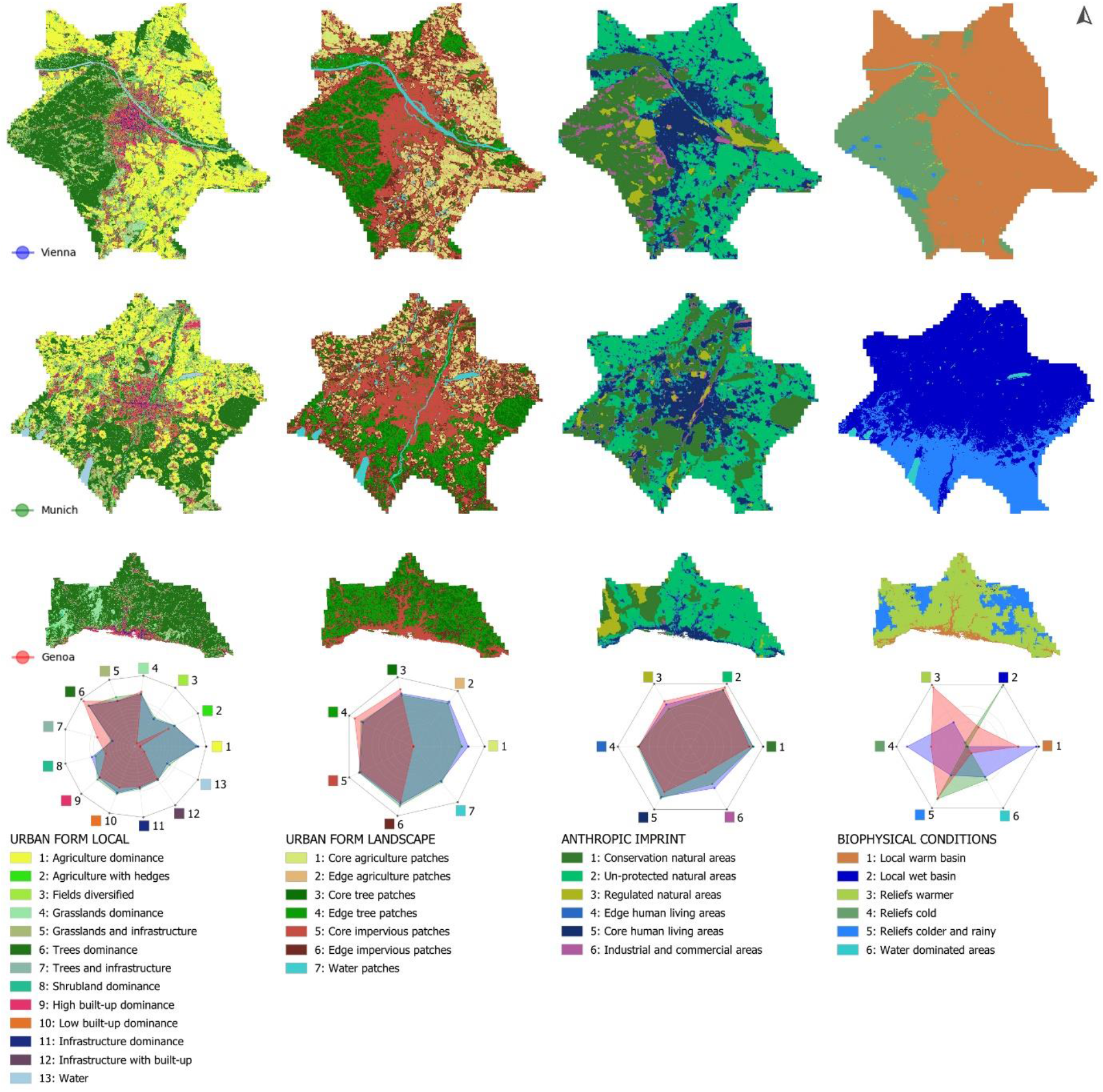
1 to 5.4: The Cohabscape methodology applied to the case studies of Vienna, Munich, and Genoa. For each sub-classification, the spread of class frequencies across the cities is shown using logarithmic scale log10. Continental cities such as Vienna and Munich share numerous Urban Form features in the peri-urban area, such as vast agricultural areas that are not present in the narrow coastal city of Genoa. On the Anthropic Imprint, the cities behave more similarly, with the only class with different values being areas with large industrial use that are less present in Genoa. Finally, the most evident sub-classification in term of regional difference is the Biophysical Conditions, indicating underlying differences even among the cities that appeared more similar from a climatic point of view (Vienna and Munich).

***The Landscape Urban Form sub-classification*** was obtained after the local scale sub-classification, using the local KPCA results as constituents. The dimensionality reduction through PCA produced six components from 24, that were grouped via Elkan KMeans algorithm into seven distinct classes (Fig. 5.2). Similarly to the local behaviour, the number of natural-dominated classes was higher (five) than those strongly influenced by artificial features (two).

***The Anthropic Imprint sub-classification*** started from reducing the number of its features from 15 to five components through PCA. Afterwards, using Elkan KMeans, six distinct classes were formed (Fig. 5.3). In this case, the subdivision was equally between landscapes dominated by anthropic use and functions, and those without heavy human presence.

Finally, ***the Biophysical Conditions sub-classification*** was formed by firstly reducing 19 features down to eight through PCA. Then, using Elkan KMeans, six classes were formed (Fig. 5.4). In this case, the resulting classes did not clearly defined a division of anthropic and natural landscapes. The delineation of urban-rural perimeter could not unambiguously be determined, and as such the classes observed primarily the most relevant regional climatic trend, together with the presence of reliefs and hills.

Each sub-classification best algorithm was selected via multiple scores observation (Annex C). The symbology and definition of class content was determined based on the PCA/KPCA values (Annex D). We plotted each map and visually confronted the results to be identify the class spread and frequency of occurrences in the different cities (Fig. 5). Notably, in for most sub-classifications, Genoa appeared as the outliner. In the case of Local and Landscape Urban Form, the continental case studies provided larger frequencies of classes correlated to agriculture and grasslands, almost not existent in Genoa.

From the Anthropic Imprint perspective, the differences between cities were minors, with Genoa possessing fewer industrial and commercial areas. The Biophysical Conditions instead provided a significantly different distribution of classes for all cities. These regional differences can be observed, as an example, in the case of urban parks (Fig. 6).

**Fig. 6:**
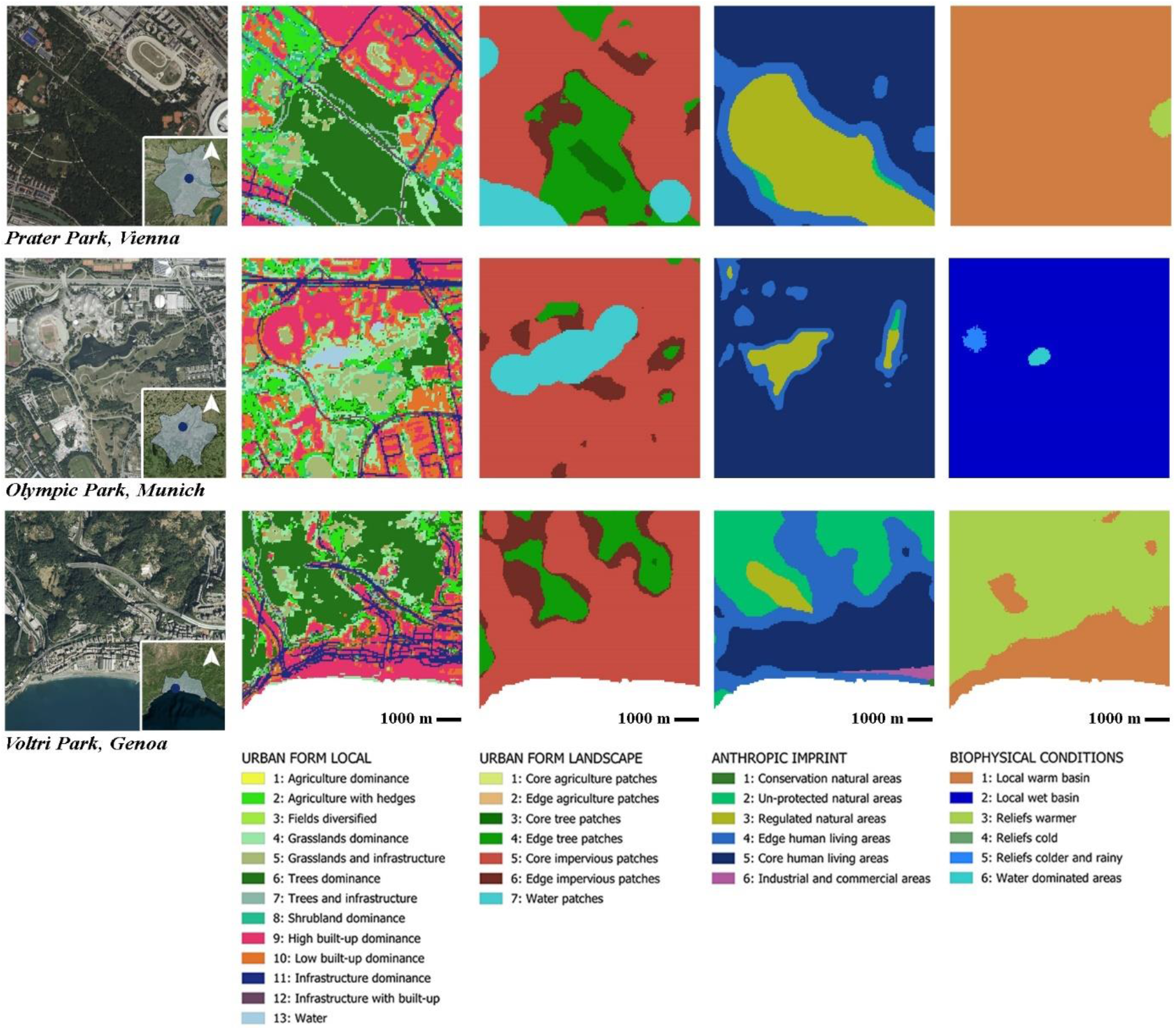
The Cohabscape methodology, zoomed-in view for three different urban parks in Vienna, Munich, and Genoa, at 1:10000 scale. This example helps understanding some of the regional differences between the different case studies, while giving an insight at the results at a finer scale. Significant differences are present in the Anthropic Imprint in terms of accessibility to Regulated natural area (class 3) in Genoa contrary to Munich and Vienna. Also, significant differences are present in the Biophysical conditions, but some insights are presents for local warmer areas (as in Genoa, where the city is highlighted primarily from temperature).

### 3.2 Comparison across different sub-classifications

We registered several key differences between the urban centre and peri-urban areas, with the former being much smaller, corresponding to 15% of the whole FUA. For simplifying the reading of our findings, we grouped each sub-classification classes together based on the dominant variable. Local Urban Form was grouped by agricultural, grasslands, trees, artificial classes, and water. Similarly, Landscape Urban Form was grouped by agricultural, trees, impervious, and water. Anthropic Imprint was divided between natural and human centric. Biophysical Conditions was divided between low and high topography, and water (Fig 7).

**Fig. 7:**
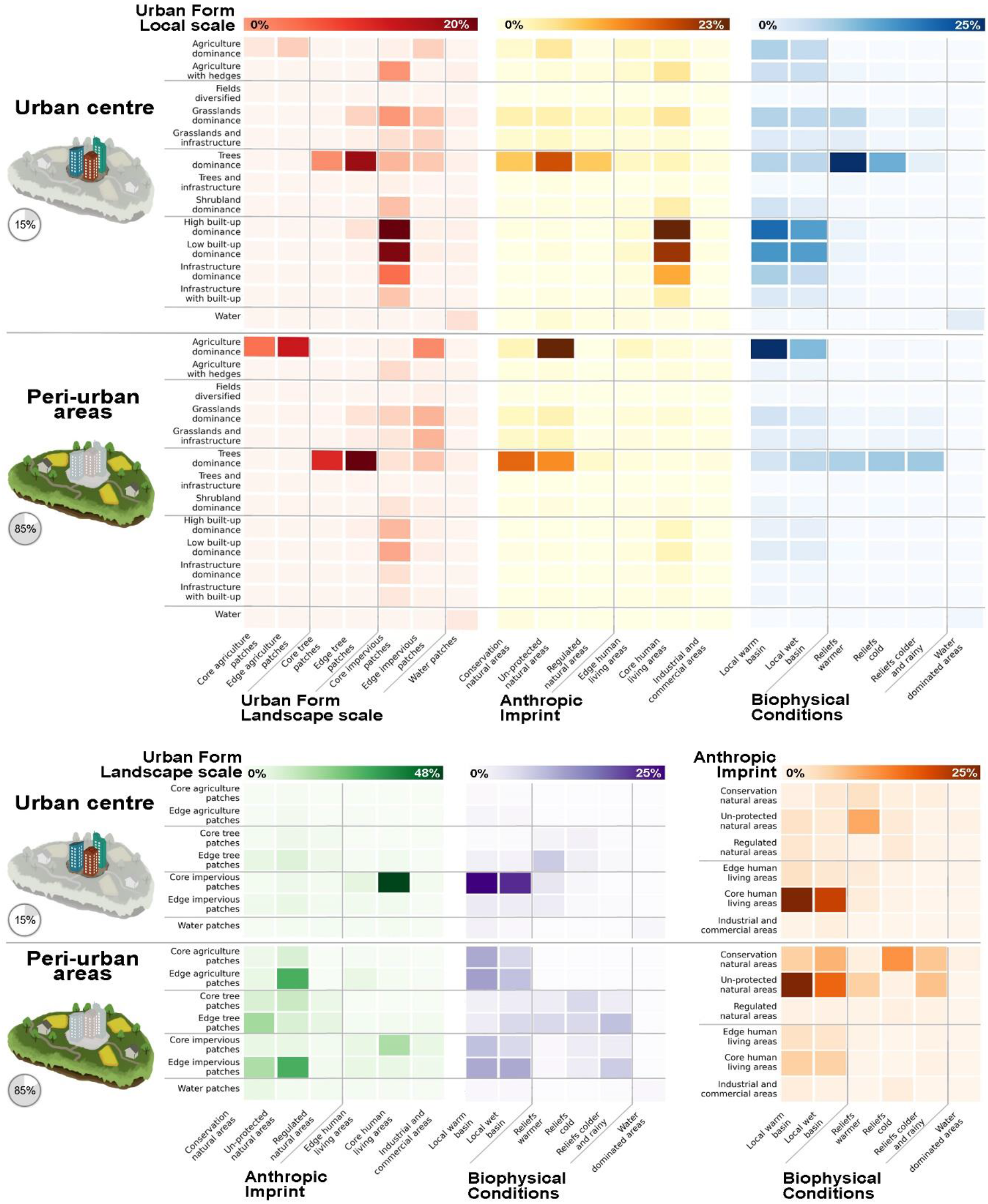
Pairwise combinations between the different sub-classifications classes frequencies of occurrences, of all case studies jointed together. Each row confronts local Urban Form with one different sub-classification between Urban Form at landscape scale, Anthropic Imprint, Biophysical Conditions. The rows are divided between the total amount in the total amount of cells in urban centres (15%), and total amount in peri-urban areas (85%).

The main correlations between urban centre and peri-urban areas highlighted different behaviours of the FUA subdivision from the different sub-classifications, primarily the different concentration of natural and artificial dominated classes, the juxtaposition of natural and impervious core and edge patches, the net human presence in terms of human-centred and natural-centred classes, and the different distribution of local temperatures and topographical features.

Generally, the urban centre conditions appeared more heterogeneous than in the peri-urban areas while looking at the Local Urban Form (Fig. 7, upper). Conversely, urban centre conditions became less heterogeneous when Local Urban Form was not a discriminating factor (Fig. 7, bottom).

## 4. Discussion

### 4.1 Urban landscapes classification to describe the diversity of urban dimensions

The proposed urban landscape classification aims to describe the diverse urban dimensions through variables important to both human and non-human inhabitants, using four separated sub-classifications. We applied the methodology to three distinct case studies and analysed the consistency and frequencies of all classes between different regions and sub-classifications. This approach highlighted numerous understandings of urban dimensions that may be applied for representing the interactions between human and non-human city inhabitants (Bettencourt, 2021). By disentangling the urban landscape description through four sub-classification, our approach aims to facilitate exploring and understanding the diversity of urban dimensions. This property corresponds to our classification’s inherent flexibility, empowering users to selectively choose and combine the four sub-classifications according to their specific needs.

### 4.2 Stratification of urban dimensions through different spatial signatures

Urban landscapes have been traditionally described as a composition of geographical and cultural attributes, such as the environment, identity, and legibility (Lynch, 1960; Wu et al., 2013); biotic attributes, namely the humans that live and interact on the landscape (Opdam et al., 2018); and the synergy between geographical, and human-induced forces (Opdam et al., 2018). Geographical attributes include the built and natural environments and have been shown to immediately shape and condition the form and texture of landscapes they are found in (Kropf, 2009). The way that humans develop their settlements influences the way in which they interact and shape the geographical attributes of the landscape (Oliveira, 2022), where climate has been shown to be the strongest driver of human development (Olgyay, 2015). In general, the preindustrial human interaction and shaping of their immediate surroundings have had low impact upon the geographical attributes (Coutard and Florentin, 2024), however, it has been reported that with the coming of the industrial age, human interaction with geographical attributes through extreme industrial activity has created the long-term effect of climate change (Bulkeley, 2010). Additionally, the passive interaction of human settlements upon the geographical attributes of the landscape leaves an urban spatial signature which is unique to the functional background of these settlements (Arribas-Bel and Fleischmann, 2022). This rendition could still take into consideration cultural attributes in the description of urban landscapes, despite the primary role of environmental characteristics. The spatial signature of human presence relevance could also be described as an hauntology, or the way humans affect the landscape even when they are not actively there (Sterling, 2022). In any case, by disentangling the urban dimensions through the use of different sub-classifications, we highlighted the possibility to confront with one or multiple dimensions and read different urban and peri-urban signatures.

### 4.3 Representing multistakeholder liveability

The traditional description of urban landscapes is conducted from the vantage point of singular types of inhabitants, predominantly human if referring to urban policies (Puchol-Salort et al., 2021; Spiliotopoulou and Roseland, 2022); or from the point of view of plants and animals for nature conservation (Ettinger et al., 2021; Klaus and Kiehl, 2021). This description of urban landscapes from the vantage point of singular user type is well-suited for their intended audience, but this approach often lack flexibility when applied to non-targeted stakeholder who perceive the environment differently; or to understand more complex dependencies of humans to non-human urban inhabitants (Likens and Lindenmayer, 2012). There are many studies that provide examples of how singular stakeholder descriptions affect the non-targeted users, with two recent examples being the creation of green infrastructure for supporting biodiversity and solutions for combating urban heat. Green infrastructure has been predominately used for fostering biodiversity and supporting numerous ecological services, however studies, such as Fairbairn et al., 2024 and Apfelbeck et al., 2020, have shown that green infrastructure positively transforms cities and improving the specific species well-being, even in cases where humans are not the targeted user. Addressing warm urban landscapes through architectural transformation to improve the quality of the human residences lives and reduce overheating caused mortalities, helps animals and plants as well (Čeplová et al., 2017). The ECOLOPES project highlights the complex challenge of considering the multistakeholder interactions in cities via the design of nature-based solutions that cater for the liveability of the different human and non-human inhabitants (Weisser et al., 2023). This is particularly challenging if the interactions are considered combined. As the paradigm for a perspective shift is often called for (Shingne and Reese, 2022; UN-Habitat, 2022), we believe that a solution for a more accurate representation of liveability could lie in the description of urban landscapes where the multiple stakeholders live and interact.

The Cohabscape methodology is designed to be coupled with city studies and comparisons of urban liveability across different regions. This classification does not serve as an omniscient knowledge base but presents a description of urban landscapes that could bridge different disciplines. The relevance of this broad and harmonised definition of urban landscapes could also be relevant for administrators and politicians. As global urbanisation continues, the population of human, plants, and animal urban inhabitants are anticipated to share an increasing area of communal spaces found within cities (Petersen et al., 2020; Rega-Brodsky et al., 2022). Consequently, urban policies need to reassess how human city dwellers can depart from the traditional vision of unsustainable development (Shingne, 2020) and modify their living norms to adapt to coexistence with non-human urban inhabitants. Our approach for comparing urban landscape diversity can help linking and strengthening the scarce and debated directives for multistakeholder cohabitation (Roudavski, 2020; European Environment Agency, 2023). This bridging approach could be enlarged to encompass other major challenges to urban liveability that have been often represented by closed-frameworks, such as climate change adaptation (Pettorelli et al., 2021; Soergel et al., 2021) and human health and well-being (Cloutier and Pfeiffer, 2015; Naeem et al., 2016).

As indicated before, the implications of such approach could be found in both green urban design, planning and ecology, when confronting with biophilic urban designs and analysis. For instance, plant species across large territories could be analysed together their capacity of helping with multi-risk climate change adaptation such as overheating phenomena and stormwater retention. Animal connectivity could be paired with our classification, by studying how optimizing home ranges could be upscaled and confronted in different locations. The use of multiple scales is useful in this regard, as it can be applied to other data, such as species occurrence and home ranges. Additionally, empirical evidence suggests that the generic and heterogeneous nature of similar descriptions of urban landscapes could outperform species-centric approaches even when explaining biodiversity patterns at the species level (Mimet et al., 2014; Hughes et al., 2021), when species-specific data is insufficient (Kattwinkel et al., 2009) or for lost taxa (Martin et al., 2023).

### 4.4 Conclusions and further developments

This work intends to present the methodology applied to a limited number of case studies. The primary outputs of this work, i.e. the four sub-classifications for the three case studies, are available in raster format. Additionally, the full extent of the PCA/KPCA dataset for all sub-classifications is available in parquet table format, allowing the comparison of the categorical class definition with the full data gradient. From this, a second, larger and more fine-tuned classification is expected to be constructed by introducing a set of improvements, i.e. larger extent, different algorithms, and revisioned data. The second step could enlarge to the full extent of the European region, considering a much higher number of FUA. This would in turn introduce a lot of computational complexity, that would be confronted by a restructuring of the algorithms used. Additionally, albeit KMeans algorithm is fast and can produce robust results with limited computation timings, hierarchical agglomeration could better represent the diversity introduced by the heterogeneous set of variables. Hierarchical agglomerations could also open up the possibility to deliver the classification at different levels of tree pruning, offering different scales of interpretation of urban landscapes. Finally, a point can be made regarding the scope of analysis could be setting the focus on just the core urban areas, over the whole FUA. Municipalities are typically the norm of landscape analysis, as most high resolution data is available and produced for its boundaries, such as temperature data and building height layers. As for the High Resolution Layers suite demonstrates, this limitation could be overcome in the foreseeable future, introducing the possibility of working on large boundaries and keeping very accurate data.

## Supporting information

Annex D

Annex A

Annex B

Annex C

## Acknowledgements

The authors gratefully acknowledge funding of the EU H2020 FET-OPEN project ECOLOPES (https://ecolopes.org/; Grant Agreement Number 964414). G.O.’s scholarship is founded by the Italian “Piano Nazionale di Ripresa e Resilienza” (D.M. n.351 09/04/2022).

This publication has been prepared using European Union’s Copernicus Land Monitoring Service information. The DOI links are located in the References section.

## Conflict of Interest

The authors declare no competing interests.

## Data availability

The code and data produced in the publication can be found respectively at www.github.com/gabrieleoneto/urban-cohabscapes and www.zenodo.org/records/14000478. Additional data, such as preliminary results, can be requested from the corresponding author.

## Author contributions: CRediT

Conceptualization: A.M., G.O., K.P., V.C., Data curation: G.O., Formal analysis: G.O., Funding acquisition: W.W., K.P., A.M., Methodology: A.M., G.O., Software: G.O., Visualization: G.O., Writing – original draft: G.O., Writing – review and editing: A.M., K.P., M.C., C.V., W.W., G.O.

## Notes

https://zenodo.org/records/14000478

https://github.com/gabrieleoneto/urban-cohabscapes

### Competing Interest Statement

The authors have declared no competing interest.

