## Supplementary material for "Urban Cohabscapes: exploring European Co-Habitative landScapes diversity, in the ECOLOPES framework": Annex D

### Annex D. Clusters definitions

Having set an optimal k for the unsupervised classification, we then need to deduce the contents of each class from its defining features. To do this, we plot the Principal Components (PC) values for each cluster using a boxplot. We rescale the components from -1 to 1 to better appreciate the higher and lower values in a comparable range, limiting spike values. We use matplotlib for all plots; we also highlighted with different colours four ranges of 0.5 from max to min to help read the graphs. We primarily focused on PC median values inside (0.3, 1) and (-1, -0.3) ranges, signifying stronger contributions and gave more importance to PC axis with thinner interquartile range.

#### Urban Form Local

Homogeneous water was not present in the PCA from the local scale, the modelling considered water as a negative presence; therefore, the patterns of water presence at the local scale is obtained by considering this negative space inside the city boundary, consisting of a final 13th cluster. Among the modelled pixels, we found 12 clusters of heterogeneous size with the minimum being 0.34% of all modelled cells (Cluster 4) and the maximum 33.44% (Cluster 3) (Fig. D.1), defined as follows based on the PCs values in Table D.1.

| Cluster | Notable PC dependencies | Description |
|---|---|---|
| 1 | Agriculture (PC1) | Agriculture dominance |
| 2 | Shrubs (PC4), agriculture (PC1) | Agriculture with hedges |
| 3 | Agriculture (PC1), grass (PC3), diversity (PC5, PC6) | Fields diversified |
| 4 | Grassland (PC2), building height (PC5, 6), built-up (PC5) | Grasslands dominance |
| 5 | Building height, streets (PC2), grasslands (PC3) | Grasslands and infrastructure |
| 6 | Trees (PC1) | Trees dominance |
| 7 | Trees (PC1), Streets and railways (PC5) | Trees and infrastructure |
| 8 | Shrubs (PC4) | Shrubland dominance |
| 9 | Built-up fraction, impervious density (PC2, PC3), building height, streets (PC3) | High built-up dominance |
| 10 | Building height, streets (PC2) | Low built-up dominance |
| 11 | Built-up fraction, impervious density (PC2, PC3), streets (PC5) | Infrastructure dominance |
| 12 | Building height (PC2, 3, 6), streets (PC2), built-up (PC3), streets (PC5) | Infrastructure with built-up |
| 13 | | Water |

Table D.1: Description of Urban Form Local classes based on PC values, highlighting dependencies based on boxplot spread and median position within the range of each component dominion.

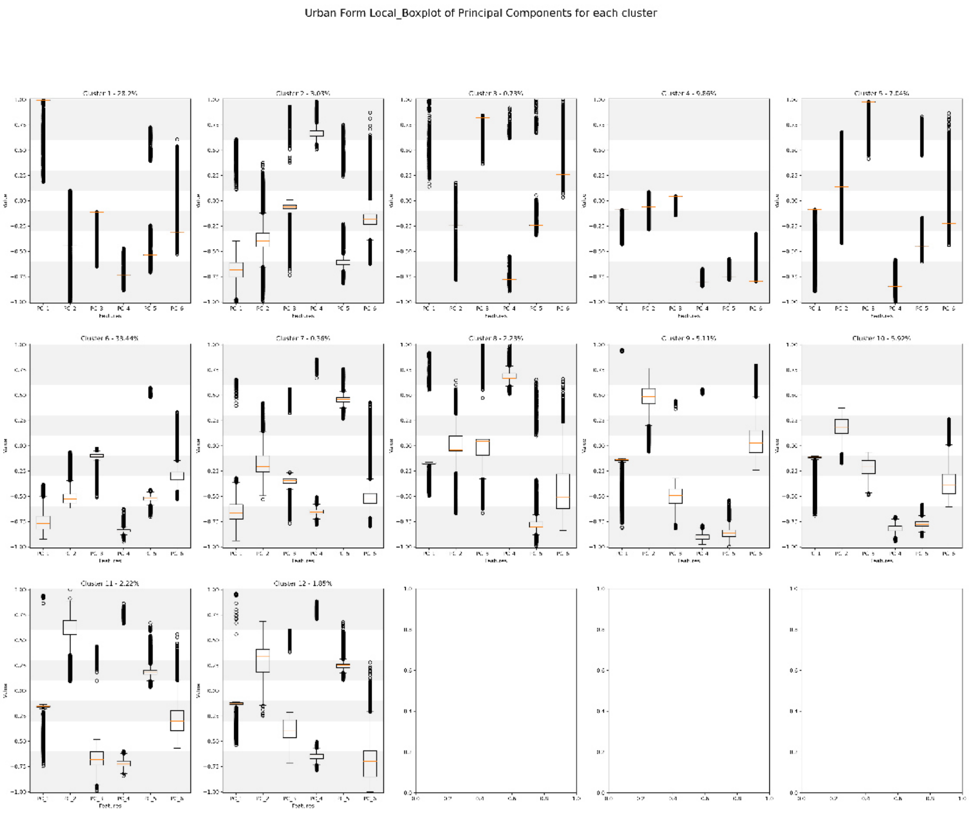

Figure D.1: Boxplot of each cluster values for Urban Form Local layer, with ranges for each corresponding Principal Component spread.

#### Urban Form Landscape

Similarly to the local scale, water was not present in the PCA; therefore, modelling considered water as a negative presence then added at the end as a final class. From the modelled values, we found 6 homogeneous clusters ranging from the smallest of 8.05% (Cluster 3) to the largest of 23.5% (Cluster 6), defined in Table D.2 based on the boxplot (Fig. D.2). We noted a trend of core clusters (3, 5, 6) with corresponding heterogeneous edge patches that came from the presence of disruptive elements (clusters 1, 2, 4).

| Cluster | Notable PC dependencies | Description |
|---|---|---|
| 1 | Contagion (PC2), std PC1 (PC2), PC2 (PC5), mean PC3 (PC5), std PC5 (PC6) PC5 (PC5), PC6 (PC5, 7), mean PC4 (PC7); most agriculture and diversity coming from aggregate areas where agriculture is present | Core agriculture patches |
| 2 | Mean PC1 (PC2), std PC2 (PC3), PC3 (PC3) | Edge agriculture patches |
| 3 | PC4 (PC3), std PC1 (PC4), std PC3 (PC4), std PC2 (PC6), std PC5 (PC6, 7), PC6 (PC7); trees and urban greenery with few disturbances | Core tree patches |
| 4 | Contagion (PC3), PC2, mean PC3, PC5, PC6 (PC5, PC6, PC7); trees with numerous disruptions coming from different sources | Edge tree patches |
| 5 | Mean PC1 (PC1), PC4 (PC1, PC3); mostly impervious and built-up areas | Core impervious patches |
| 6 | Mean PC3 (PC1, PC2), mean PC4 (PC2,7), std PC2 (PC3, 6), PC3 (PC3), std PC1 (PC4), std PC3 (PC5), std PC5 (PC6, 7), PC6 (PC7) | Edge impervious patches |
| 7 | | Water patches |

Table D.2: Description of Urban Form Landscape classes based on PC values, highlighting dependencies based on boxplot spread and median position within the range of each component dominion. The dependencies refer to local mean and std PC axis, keeping in parentheses the corresponding landscape components. Additional notes are added to help interpretation.

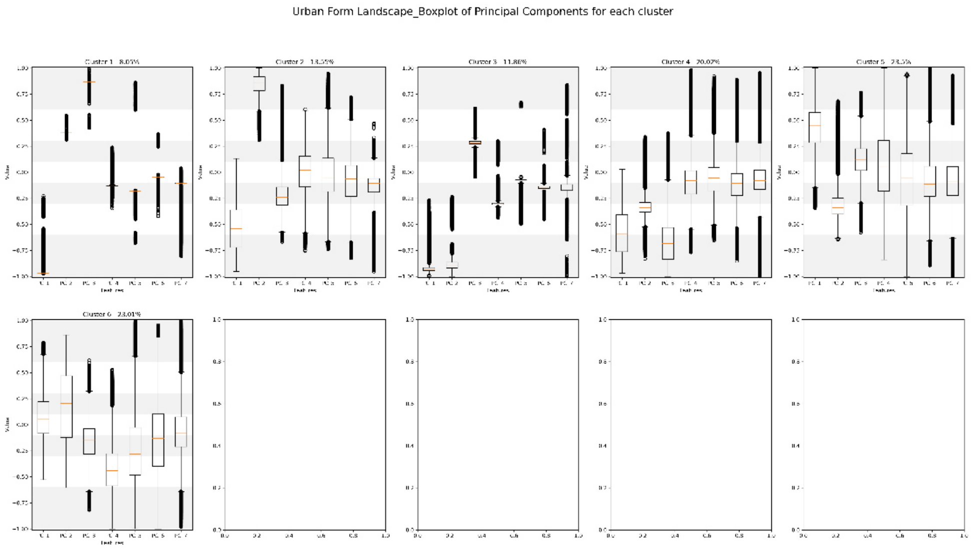


Figure D.2: Boxplot of each cluster values for Urban Form Landscape layer, with ranges for each corresponding Principal Component spread.

#### Anthropic Imprint

We modelled the entire collection of pixels, without removing classes before preprocessing. From the modelled values, we found 6 very inhomogeneous clusters ranging from the smallest of 3.13% (Cluster 3) to the largest of 42.08% (Cluster 1), defined in Table D.3 based on the boxplots (Fig. D.3). We noted

| Cluster | Notable PC dependencies | Description |
|---|---|---|
| 1 | Public green (PC1), commercial area (PC1), industrial area (PC1), public service (PC1), preservation policy (PC2) | Conservation natural areas |
| 2 | Age groups (PC2), public green (PC3), recreation area (PC2, 4), transports (PC4); places without human presence and no significant services, but very high distance to public transports | Un-protected natural areas |
| 3 | Population density (PC1), age groups (PC1), | Regulated natural areas |
| 4 | Commercial area (PC1), industrial area (PC1), public service (PC1), transports (PC4) | Edge human living areas |
| 5 | Age groups (PC2), recreation area (PC2) | Core human living areas |
| 6 | Commercial area (PC1), industrial area (PC1), public service (PC1); dense industrial and commercial activities | Industrial and commercial areas |

Table D.3: Description of Anthropic Imprint classes based on PC values, highlighting dependencies based on boxplot spread and median position within the range of each component dominion.

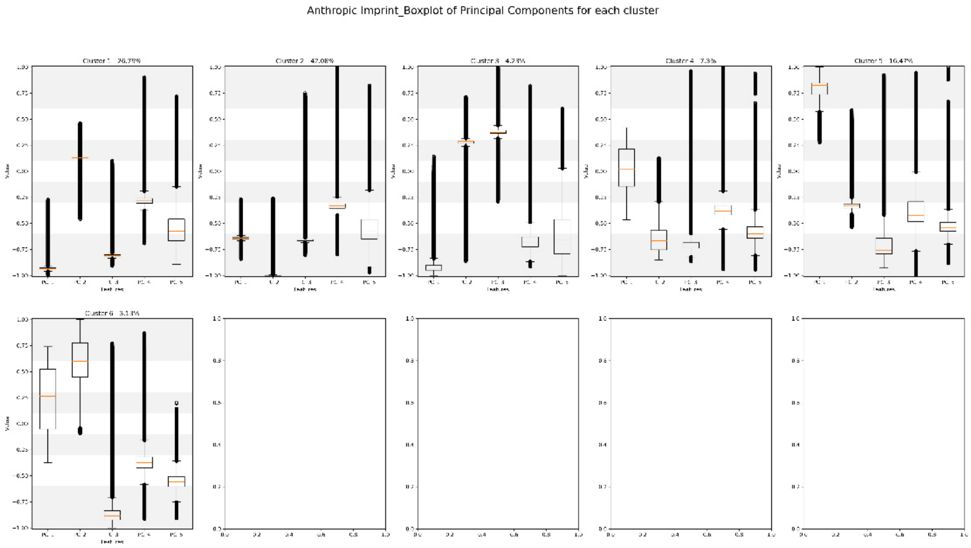


Figure D.3: Boxplot of each cluster values for Anthropic Imprint layer, with ranges for each corresponding Principal Component spread.

#### Biophysical Conditions

We modelled the entire collection of pixels, without removing classes before preprocessing. From the modelled values, we found 6 clusters ranging from the smallest of 7.53% (Cluster 6) to the largest of 33.32% (Cluster 2), defined in Table D.4 based on the boxplots (Fig. D.4). The regional difference posed significant difficulty in defining the discrepancies between clusters. Similarly with differentiating between temperature behaviour and soil constitution. As such, this layer was the only that produced some clusters that could not be found in specific cities: Cluster 3 is absent from Munich and Cluster 2 is absent from Vienna.

| Cluster | Notable PC dependencies | Description |
|---|---|---|
| 1 | Temperature max and min relative (PC1), wind speed (PC1), precipitations absolute (PC8) | Local warm basin |
| 2 | Altitude std (PC1, 2), slope mean (PC1, 2, 3) and std (PC3), aspect mean (PC1), curvature std (PC1, 2, 3), precipitations absolute (PC8) | Local wet basin |
| 3 | Altitude std (PC1), slope (PC1, 3), aspect mean (PC1, 3) and std (PC6), curvature std (PC1, 3), temperature max and min absolute (PC2, 3) | Reliefs warmer |
| 4 | Wind speed (PC1, 4), soil wetness (PC2), temperature max and min relative (PC2), precipitations absolute (-) | Reliefs cold |
| 5 | Precipitations relative (PC1, 2) and absolute (PC2, 8), slope mean (PC3), curvature std (PC3), wind speed (PC4) | Reliefs colder and rainy |
| 6 | Wind speed (PC1), permanent and temporary water (PC4, 5), temperature max and min relative (PC4), precipitations absolute (PC8) | Water dominated areas |

Table D.4: Description of Biophysical Conditions classes based on PC values, highlighting dependencies based on boxplot spread and median position within the range of each component dominion. Under PC dependencies, some comments were added to improve understanding.

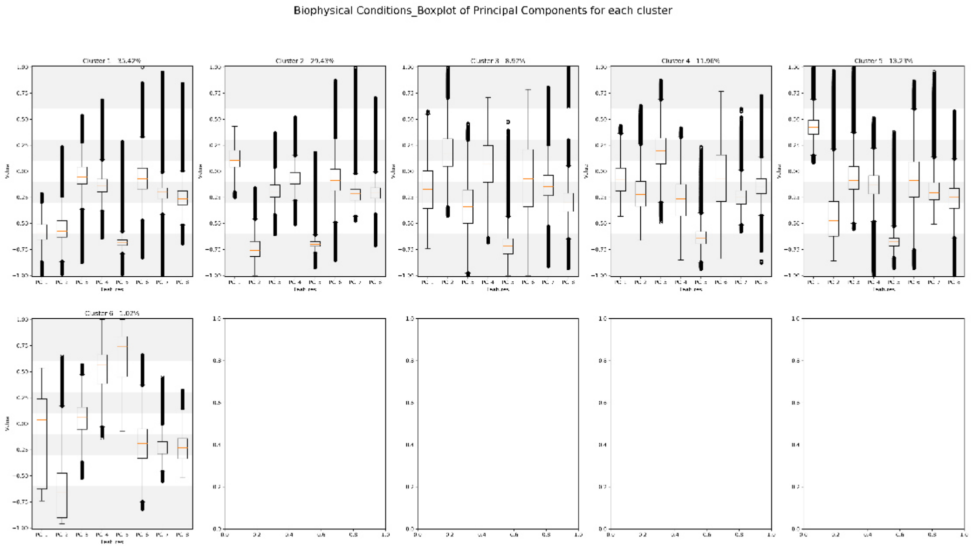


Figure D.4: Boxplot of each cluster values for Biophysical Conditions layer, with ranges for each corresponding Principal Component spread.
