## Supplementary material for "Urban Cohabscapes: exploring European Co-Habitative landScapes diversity, in the ECOLOPES framework": Annex A

### Annex A. Variable tables, methodologies and data origin

In this study, we chose an operative threshold of what is considered ‘urban’ that could overcome the immediate human-centric perspective of city borders. This could be the case for administrative boundaries. We choose to adopt the Global Human Settlement Layer definition of Functional Urban Areas (Schiavina et al., 2023) as they represent urban areas in a broader way, considering both urban centres and peri-urban areas that economically and infrastructurally contribute in defining and shaping cities.

We chose an equiareal projection to force all the grid cells in our modelling to represent the same spatial extent, therefore comparing urban landscapes of the same size. Moreover, each city was aligned on the grid by locating the corresponding closest points. The urban boundaries were buffered by 200-metres, adjusting for possible edge effects that could had arisen during computations. Each urban boundary was then rasterised and used as a reference raster for all further operation, such as matching and stacking.

The datasets were manually downloaded, but automatic scraping was conducted where relevant or meaningful. The OSMNX package provided a flexible tool for extracting OSM geometry based on tags (Boeing 2024). We used this approach for each OSM-based variable, selecting a roster of OSM tags and accurately pruning and cleaning the geometries. Since OSM feature tagging can be unreliable, we opted for a “all-or-nothing” approach, were we firstly disregarded too specific tags, and then selected the most popular areas and places tags. We only kept geometries and rasterized them onto a grid, keeping information as only presences. For other datasets, custom API were adopted were available, helping with the handling of large datasets and selecting features, (notably, the chelsa_cmip6 package; Karger et al., 2017).

Since most datasets were distributed in a tile format, we considered constructing mosaics for each one. This would have produced very large and memory intensive rasters and complicated the computation process. Instead, the downloaded data was extracted and read with a feature search approach tile-by-tile, scanning each tile for information within the city boundary mask, and mosaicking the fragments only after. We had to read the first tile CRS and reproject our mask to it, then revert the data back to our Mollweide grid and assuring each pixel received information. This method benefits the scalability of our methodologies by optimizing memory usage.

The computation pipeline was structured in two different procedures, depending on the variable being at local scale or at landscape scale. The former is considered only for the Urban Form local sub-classification, being the one with native resolutions of 10-meter or finer. The latter approach is considered for remaining sub-classifications, as most variables have coarser resolutions that would be better computed at a higher scale. We run landscape metrics through a rounded 21 cell-diameter window, operating both kernel averages and standard deviations. Then we stacked the resulting data onto the reference raster, reprojecting to Mollweide, and synching resolution by resampling at 10-meter. Most of the numeric computation was conducted on xarray DataArrays; the vectorial information was handled by geopandas and the raster writing was conducted using rasterio and rioxarray. Kernel calculations and distances were computed using scipy.

| **Urban Form Local** | | | | |
|---|---|---|---|---|
| **Category** | **Variable** | **Methodology** | **Data origin** | **Native resolution** |
| **Artificial features** | Building Height | Both sources are reprojected and matched to the reference grid using nearest neighbours. Then, UA BH 2012 layer is superimposed over MSZ Classes of building height (3<, 3-6, 6-15,15-30,>30). | Global Human Settlement Layer / Built Classification with Morphological Settlement Zones (MSZ) 10.2905/3C60DDF6-0586-4190-854B-F6AA0EDC2A30 + Copernicus / Urban Atlas Building Height 2012 https://doi.org/10.2909/42690e05-edf4-43fc-8020-33e130f62023 | 10 |
| | Built-up Fraction | The source is reprojected and matched to the reference grid using nearest neighbours. | Global Human Settlement Layer / Built Surface (Built-s) 10.2905/9F06F36F-4B11-47EC-ABB0-4F8B7B1D72EA | 10 |
| | Mayor Transport route Presence | OSM geometries are extracted using osmnx with the base grid as a mask. The OSM geometries are then rasterized matching the reference grid, and added together with “street” layer from MSZ classification. The data is then reprojected matching the source grid. | Open Street Maps (OSM) / osm_tags = [{'highway':"motorway"},<br>{'highway':"motorway_link"},<br>{'highway':"trunk"},<br>{'highway':"trunk_link"},<br>{'highway':"primary"}, | 10 |

| | | | | |
|---|---|---|---|---|
| | | | {'highway':"primary_link"},<br>{'highway':"secondary"},<br>{'highway':"secondary_link"},<br>{'highway':"tertiary"},<br>{'highway':"tertiary_link"},<br>{'highway':"unclassified"},<br>{'highway':"road"},<br>{'highway':"residential"},<br>{'railway':True}<br>];<br>Global Human Settlement Layer / Built Classification with Morphological Settlement Zones (MSZ) 10.2905/3C60DDF6-0586-4190-854B-F6AA0EDC2A30 | |
| | Imperviousness Density | The source is reprojected and matched to the reference grid using nearest neighbours. | Copernicus High Resolution Layer / Imperviousness https://doi.org/10.2909/3bf542bd-eebd-4d73-b53c-a0243f2ed862 | 10 |
| **Natural features** | Tree Cover Density | The source is reprojected and matched to the reference grid using nearest neighbours. | Copernicus High Resolution Layer / Tree Cover Density https://doi.org/10.2909/486f77da-d605-423e-93a9-680760ab6791 | 10 |
| | Grassland Presence | The source is reprojected and matched to the reference grid using nearest neighbours. | Copernicus High Resolution Layer / Grassland https://doi.org/10.2909/60639d5b-9164-4135-ae93-fb4132bb6d83 | 10 |
| | Shrubs Presence | The source is reprojected and matched to the reference grid using nearest neighbours. | Copernicus High Resolution Layer / Small Woody Features https://doi.org/10.2909/a8e683b1-2f96-45c8-827f-580a79413018 | 5 |
| | Agricultural Land Presence | The source layer "crop" is reprojected and matched to the reference grid using nearest neighbours. | European Space Agency / WorldCover V2 2021 https://doi.org/10.5281/zenodo.7254221 | 10 |
| | Aquatic Surface Presence | The source layer "permanent water" is reprojected and matched to the reference grid using nearest neighbours. | Copernicus High Resolution Layer / Water and Wetness Water and Wetness status 2018 (raster 10 m and 100 m), Europe, 3-yearly — Copernicus Land Monitoring Service | 10 |
| | Diversity of land cover in cell | Sum of land covers in cell: built-up fraction, transports, trees, grass, shrubs, agriculture | | 10 |

| **Urban Form Landscape** | | | | |
|---|---|---|---|---|
| **Category** | **Variable** | **Methodology** | **Data origin** | **Native resolution** |
| **Class metrics** | Mean and standard deviation of each Urban Form Local Principal Component | Mean and standard deviation computed using a 21x21 (100m radius) circular kernel with scipy generic filter. | Urban Form Local classification as base mosaic | 10 |
| **Landscape metrics** | Shannon Diversity Index | Using Fragstats 4.2, computed in a 100m radius rounded window | Urban Form Local classification as base mosaic | 10 |
| | Contagion | Using Fragstats 4.2, computed in a 100m radius rounded window | Urban Form Local classification as base mosaic | 10 |

| **Anthropic Imprint** |
|---|

| Category | Variable | Methodology | Data origin | Native resolution |
|---|---|---|---|---|
| Demographics | Population Density | The source is reprojected and matched to the reference grid using nearest neighbours. Then, the average is computed in a moving window of nearest neighbour resampled value of raster; using scipy generic filter as a 21x21 round kernel (100m diameter). | Global Human Settlement Layer (GHSL)/Population Density (POP) 10.2905/2FF68A52-5B5B-4A22-8F40-C41DA8332CFE | 100 |
| | Age Structures: 0-15 | The source is divided in age groups and unified by merging female and male layers, each opportunately reprojected and matched to the reference grid, using the nearest neighbours. Then, a ratio is computed by dividing the 0-15 partition with the sum of all layers. Finally, the average is computed in a moving window using scipy generic filter as a 21x21 round kernel (100m diameter). | WorldPop / Age and gender structures, structured 10.5258/SOTON/WP00695 | 100 |
| | Age Structures: 20-35 | The source is divided in age groups and unified by merging female and male layers, each opportunately reprojected and matched to the reference grid, using the nearest neighbours. Then, a ratio is computed by dividing the 20-35 partition with the sum of all layers. Finally, the average is computed in a moving window using scipy generic filter as a 21x21 round kernel (100m diameter). | WorldPop / Age and gender structures, structured 10.5258/SOTON/WP00695 | 100 |
| | Age Structures: 40-60 | The source is divided in age groups and unified by merging female and male layers, each opportunately reprojected and matched to the reference grid, using the nearest neighbours. Then, a ratio is computed by dividing the 40-60 partition with the sum of all layers. Finally, the average is computed in a moving window using scipy generic filter as a 21x21 round kernel (100m diameter). | WorldPop / Age and gender structures, structured 10.5258/SOTON/WP00695 | 100 |
| | Age Structures: 65-85 | The source is divided in age groups and unified by merging female and male layers, each opportunately reprojected and matched to the reference grid, using the nearest neighbours. Then, a ratio is computed by dividing the 65-85 partition with the sum of all layers. Finally, the average is computed in a moving window using scipy generic filter as a 21x21 round kernel (100m diameter). | WorldPop / Age and gender structures, structured 10.5258/SOTON/WP00695 | 100 |
| Land use characteristics | Commercial Area Ratio | The geometries are extracted using osmnx by using the base grid as a mask. The mask is reprojected to epsg:4326 to comply with OSM CRS, then the geometries are reprojected back to Mollweide before modelling. The geometries are then rasterized matching the reference grid. Finally, the ratio is computed by summing pixels with values of commercial buildings dividing by the total areas of the moving window, computed using scipy generic filter as a 21x21 round kernel (100m diameter). | osm_tags = [{'landuse':'commercial'},<br>{'landuse':'retail'},<br>{'craft':True},<br>{'shop':True},<br>] | Vectorial |
| | Industrial Area Ratio | The geometries are extracted using osmnx by using the base grid as a mask. The mask is reprojected to epsg:4326 to comply with OSM CRS, then the geometries are reprojected back to Mollweide before modelling. The geometries are then rasterized matching the reference grid. Finally, the ratio is computed by summing pixels with values of | osm_tags = [{'landuse':'industrial'},<br>{'landuse':'depot'},<br>{'landuse':'garages'},<br>{'man_made':'works'},<br>{'man_made':'beacon'}, | Vectorial |

| | | | | |
|---|---|---|---|---|
| | | industrial buildings dividing by the total areas of the moving window, computed using scipy generic filter as a 21x21 round kernel (100m diameter). | {'man_made':'bunker_silo'},<br>{'man_made':'chimney'},<br>{'man_made':'communications_tower'},<br>{'man_made':'crane'},<br>{'man_made':'gasometer'},<br>{'man_made':'goods_conveyor'},<br>{'man_made':'kiln'},<br>{'man_made':'lighthouse'},<br>{'man_made':'mast'},<br>{'man_made':'mineshaft'},<br>{'man_made':'monitoring_station'},<br>{'man_made':'observatory'},<br>{'man_made':'pumping_station'},<br>{'man_made':'pipeline'},<br>{'man_made':'reservoir_covered'},<br>{'man_made':'silo'},<br>{'man_made':'storage_tank'},<br>{'man_made':'tailings_pond'},<br>{'man_made':'tower'},<br>{'man_made':'wastewater_plant'},<br>{'man_made':'watermill'},<br>{'man_made':'water_tower'},<br>{'man_made':'windmill'},<br>] | |
| | Public Services Area Ratio | The geometries are extracted using osmnx by using the base grid as a mask. The mask is reprojected to epsg:4326 to comply with OSM CRS, then the geometries are reprojected back to Mollweide before modelling. The geometries are then rasterized matching the reference grid. Finally, the ratio is computed by summing pixels with values of public service buildings dividing by the total areas of the moving window, computed using scipy generic filter as a 21x21 round kernel (100m diameter). | osm_tags = [<br>{'landuse':'institutional'},<br>{'landuse':'educational'},<br># education<br>{'amenity':'college'},<br>{'amenity':'dancing_school'},<br>{'amenity':'driving_school'},<br>{'amenity':'first_aid_school'},<br>{'amenity':'kindergarten'},<br>{'amenity':'language_school'},<br>{'amenity':'library'},<br>{'amenity':'surf_school'},<br>{'amenity':'toy_library'},<br>{'amenity':'research_institute'},<br>{'amenity':'training'},<br>{'amenity':'music_school'},<br>{'amenity':'school'},<br>{'amenity':'traffic_park'},<br>{'amenity':'university'}, | Vectorial |

| | | | | |
|---|---|---|---|---|
| | | | # financial<br>{'amenity':'atm'},<br>{'amenity':'payment_terminal'},<br>{'amenity':'bank'},<br>{'amenity':'bureau_de_change'},<br>{'amenity':'money_transfer'},<br>{'amenity':'payment_centre'},<br># healthcare<br>{'amenity':'baby_hatch'},<br>{'amenity':'clinic'},<br>{'amenity':'dentist'},<br>{'amenity':'doctors'},<br>{'amenity':'hospital'},<br>{'amenity':'nursing_home'},<br>{'amenity':'pharmacy'},<br>{'amenity':'social_facility'},<br>{'amenity':'veterinary'},<br># public service<br>{'amenity':'courthouse'},<br>{'amenity':'fire_station'},<br>{'amenity':'police'},<br>{'amenity':'post_box'},<br>{'amenity':'post_depot'},<br>{'amenity':'post_office'},<br>{'amenity':'prison'},<br>{'amenity':'ranger_station'},<br>{'amenity':'townhall'},<br>] | |
| | Recreational Area Ratio | The geometries are extracted using osmnx by using the base grid as a mask. The mask is reprojected to epsg:4326 to comply with OSM CRS, then the geometries are reprojected back to Mollweide before modelling. The geometries are then rasterized matching the reference grid. Finally, the ratio is computed by summing pixels with values of recreational areas dividing by the total areas of the moving window, computed using scipy generic filter as a 21x21 round kernel (100m diameter). | osm_tags = [<br>{'landuse':'recreation_ground'},<br>{'landuse':'religious'},<br>{'leisure':True},<br>{'sport':True},<br># sustenance<br>{'amenity':'bar'},<br>{'amenity':'biergarten'},<br>{'amenity':'cafe'},<br>{'amenity':'fast_food'},<br>{'amenity':'food_court'},<br>{'amenity':'ice_cream'},<br>{'amenity':'pub'},<br>{'amenity':'restaurant'},<br># entertainment | Vectorial |

| | | | | |
|---|---|---|---|---|
| | | | {'amenity':'arts_centre'},<br>{'amenity':'brothel'},<br>{'amenity':'casino'},<br>{'amenity':'cinema'},<br>{'amenity':'community_centre'},<br>{'amenity':'conference_centre'},<br>{'amenity':'events_venue'},<br>{'amenity':'exhibition_centre'},<br>{'amenity':'fountain'},<br>{'amenity':'gambling'},<br>{'amenity':'love_hotel'},<br>{'amenity':'music_venue'},<br>{'amenity':'nightclub'},<br>{'amenity':'planetarium'},<br>{'amenity':'public_bookcase'},<br>{'amenity':'social_centre'},<br>{'amenity':'stripclub'},<br>{'amenity':'studio'},<br>{'amenity':'swingerclub'},<br>{'amenity':'theatre'},<br>] | |
| | Cultural Location Ratio | The geometries are extracted using osmnx by using the base grid as a mask. The mask is reprojected to epsg:4326 to comply with OSM CRS, then the geometries are reprojected back to Mollweide before modelling. The geometries are then rasterized matching the reference grid. Finally, the ratio is computed by summing pixels with values of cultural locations dividing by the total areas of the moving window, computed using scipy generic filter as a 21x21 round kernel (100m diameter). | osm_tags = [{'historic':True},<br>{'tourism':"alpine_hut"},<br>{'tourism':"aquarium"},<br>{'tourism':"artwork"},<br>{'tourism':"attraction"},<br>{'tourism':"gallery"},<br>{'tourism':"museum"},<br>{'tourism':"yes"},<br>{'heritage':True},<br>] | Vectorial |
| | Public Green Proportion | The geometries are extracted using osmnx by using the base grid as a mask. The mask is reprojected to epsg:4326 to comply with OSM CRS, then the geometries are reprojected back to Mollweide before modelling. The geometries are then rasterized matching the reference grid. Then, presence values (as in, Boolean 1/0) from Grass, Shrubs, and Trees (same as the local layers) are summed together to create the total amount of green. The public green is produced firstly by using the total amount of green as a mask for the OSM geometries, then the proportion is computed by dividing the remaining osm green pixels by the total amount of green in the moving window, computed using scipy generic filter as a 21x21 round kernel (100m diameter). | osm_tags = [<br>{'leisure':"park"},<br>{'leisure':"garden"},<br>{'leisure':"common"},<br>{'leisure':"playground"},<br>] | Vectorial |

| | | | | |
|---|---|---|---|---|
| | Preservation Policy Ratio | The source vector geometries are rasterized on the base grid and reprojected to Mollweide CRS (esri 54009). Then, the ratio is calculated by using a moving window, computed using scipy generic filter as a 21x21 round kernel (100m diameter), by dividing the source data by the total area of the window. | World Database of Protected Areas (WDPA)/Other Effective Area-Based Conservation Measures (WDOECM) and International Union for Conservation of Nature (IUCN) WDPA_WDOECM_Manual_1_6.pdf | Vectorial |
| | Average Distance to Public Transport | The OSM geometry is extracted using osmnx by masking with the base grid. The grid is projected to epsg 4326 to comply with OSM native CRS, then the geometries are reprojected back to Mollweide. The data is rasterized matching the reference grid. Then, for each pixel of public transport, the distance matrix is computed using scipy ndimage.distance_transform_edt sampling [10,10], equal to the reference grid. Having the distance matrix, the final average is computed by moving window using scipy generic filter as a 21x21 round kernel (100m diameter). | osm_tags = [ {'public_transport':True} ] | Vectorial |
| **Anthropic emissions** | Nighttime light Pollution | The source raster is reprojected matching the reference grid using nearest neighbours. | Wuhan University / Luojia 1-A Nighttime Lights http://59.175.109.173:8888/app/login_en.html | 130 |
| | Anthropogenic Heat Flux | The source raster is reprojected matching the reference grid using nearest neighbours. | Varquez et al. (2021)/Average Anthropic Heat Emission (AHE) 2010 10.1038/s41597-021-00850-w | 1000 |

| **Biophysical Conditions** | | | | |
|---|---|---|---|---|
| **Category** | **Variable** | **Methodology** | **Data origin** | **Native resolution** |
| **Hydrology** | Permanent water ratio | The source layer "permanent water" is reprojected and matched to the reference grid using nearest neighbours. Then, the ratio is computed using a moving window with scipy generic filter as a 21x21 round kernel (100m diameter), dividing the source data by the total area of the window. | Copernicus High Resolution Layer / Water and Wetness Water and Wetness status 2018 (raster 10 m and 100 m), Europe, 3-yearly — Copernicus Land Monitoring Service | 10 |
| | Temporary water ratio | The source layer "temporary water" is reprojected and matched to the reference grid using nearest neighbours. Then, the ratio is computed using a moving window with scipy generic filter as a 21x21 round kernel (100m diameter), dividing the source data by the total area of the window. | As before | 10 |
| **Topography** | Mean Altitude | The source DEM is reprojected and matched to the reference grid using nearest neighbours. The mean is computed using a moving window with scipy generic filter as a 21x21 round kernel (100m diameter). | EOC Geo service / TanDEM-X 30m Edited Digital Elevation Model (EDEM) https://download.geoservice.dlr.de/TDM30_EDEM/ | 30 |
| | Standard deviation Altitude | As before, but the filter computes the standard deviation | As before | 30 |
| | Mean Slope | The source DEM is reprojected and matched to the reference grid using nearest neighbours. Then, using xdem terrain.get_terrain_attribute, the slope is computed. The mean is | EOC Geo service / TanDEM-X 30m Edited Digital Elevation Model (EDEM) https://download.geoservice.dlr.de/TDM30_EDEM/ | 30 |

| | | | | |
|---|---|---|---|---|
| | | computed using a moving window with scipy generic filter as a 21x21 round kernel (100m diameter). | | |
| | Standard deviation Slope | As before, but the filter computes the standard deviation | As before | 30 |
| | Mean Aspect | The source DEM is reprojected and matched to the reference grid using nearest neighbours. Then, using xdem terrain.get_terrain_attribute, the aspect is computed. The mean is computed using a moving window with scipy generic filter as a 21x21 round kernel (100m diameter). | EOC Geo service / TanDEM-X 30m Edited Digital Elevation Model (EDEM) https://download.geoservice.dlr.de/TDM30_EDEM/ | 30 |
| | Standard deviation Aspect | As before, but the filter computes the standard deviation | As before | 30 |
| | Mean Curvature | The source DEM is reprojected and matched to the reference grid using nearest neighbours. Then, using xdem terrain.get_terrain_attribute, the curvature is computed. The mean is computed using a moving window with scipy generic filter as a 21x21 round kernel (100m diameter). | EOC Geo service / TanDEM-X 30m Edited Digital Elevation Model (EDEM) https://download.geoservice.dlr.de/TDM30_EDEM/ | 30 |
| | Standard deviation Curvature | As before, but the filter computes the standard deviation | As before | 30 |
| **Soil** | Soil wetness ratio | The source layers "permanent and temporary wetness" are reprojected and matched to the reference grid using nearest neighbours. Then, the ratio is computed using a moving window with scipy generic filter as a 21x21 round kernel (100m diameter), dividing the source data by the total area of the window. | Copernicus High Resolution Layer / Water and Wetness Water and Wetness status 2018 (raster 10 m and 100 m), Europe, 3-yearly — Copernicus Land Monitoring Service | 10 |
| **Climate** | Absolute max of air temperature, mean value | Chelsa cmip6 is used to extract data for a specific period. From the variable bio5, I extract the scenarios ssp126, ssp370, and ssp585 and average the results for Average Warmest Month Air Temperature in the period 2014-2023. Then, the data is reprojected matching the reference grid. The mean is computed using a moving window with scipy generic filter as a 21x21 round kernel (100m diameter). | CHELSA BIOCLIM + v2.1 Bioclim – Chelsa Climate (chelsa-climate.org) | 1000 |
| | Relative max of air temperature, mean value | From the result above, the relative is computed by dividing every pixel by the raster mean value. | As above | 1000 |
| | Absolute min of air temperature, mean value | Chelsa cmip6 is used to extract data for a specific period. From the variable bio6, I extract the scenarios ssp126, ssp370, and ssp585 and average the results for Average Coldest Month Air Temperature in the period 2014-2023. Then, the data is reprojected matching the reference grid. The mean is computed using a moving window with scipy generic filter as a 21x21 round kernel (100m diameter). | CHELSA BIOCLIM + v2.1 Bioclim – Chelsa Climate (chelsa-climate.org) | 1000 |
| | Relative min of air temperature, mean value | From the result above, the relative is computed by dividing every pixel by the raster mean value. | As above | 1000 |
| | Absolute precipitations, mean value | Chelsa cmip6 is used to extract data for a specific period. From the variable bio12, I extract the scenarios ssp126, ssp370, and ssp585 and average the results for Average Annual Precipitations in the period 2014-2023. Then, the data is reprojected matching the | CHELSA BIOCLIM + v2.1 Bioclim – Chelsa Climate (chelsa-climate.org) | |

| | | | | |
|---|---|---|---|---|
| | | reference grid. The mean is computed using a moving window with scipy generic filter as a 21x21 round kernel (100m diameter). | | |
| | Relative precipitations, mean value | From the result above, the relative is computed by dividing every pixel by the raster mean value. | As above | 1000 |
| | Wind speed at 10m, mean value | The source raster is extracted using the reference grid as mask, then reprojected matching the grid. Then, the mean is computed using a moving window with scipy generic filter as a 21x21 round kernel (100m diameter). | Global Wind Atlas / Wind speed 10m https://globalwindatlas.info/en/ | 300 |
| | Wind speed at 200m, mean value | The source raster is extracted using the reference grid as mask, then reprojected matching the grid. Then, the mean is computed using a moving window with scipy generic filter as a 21x21 round kernel (100m diameter). | Global Wind Atlas / Wind speed 200m https://globalwindatlas.info/en/ | 300 |
