## Supplementary material for "Urban Cohabscapes: exploring European Co-Habitative landScapes diversity, in the ECOLOPES framework": Annex B

### Annex B. Principal Component Analysis (PCA) for the four layers

We set a 0.95 expected variance for our Principal Components (PC) to explain. We deduced the minimum number of PC from the minimum number of eigenvalues necessary to explain the set expected variance. We aimed for a high expected variance to consider even minor effects from lesser variables. We compared PCA with Kernel Principal Component Analysis (KPCA), approximating kernels with the nystroem method. We used sklearn implementations for PCA, nystroem, and gridsearchcv to compute the analysis and the optimization. The resulting confront matrixes shows the covariance between the original features and the PC axes, with values from 1 (strongly positively correlated) to -1 (strongly negatively correlated). The matrixes were plotted using matplotlib and seaborn heatmap function.

#### Urban Form Local

The analysis confronted the PCA and KPCA methods with similar results. The former explained 0.95 variance with 6 PCs (Fig. B.1), while the best kernel of the latter explained 0.98 variance with the same number of PC (Fig. B.2). Therefore, we went for the KPCA set of components, as indicated from the confront plot (Fig. B.3). Notably, the kernel behaviour was linear, albeit with different hyperparameters and coming from an approximation of 50 features. We define the 6 PCs as follows in Table B.1, ignoring contributions with covariance below 0.1:

| Principal Component | Strong correlation (\|cov\| > 0.6) | Medium correlation (0.3 < \|cov\| <= 0.6) | Weak correlation (0.1 < \|cov\| <= 0.3) |
|---|---|---|---|
| PC1 | tree cover density (-), agriculture presence (+) | | |
| PC2 | built up fraction (+), impervious density (+) | building height (+), traffic routes (+), trees (-), agriculture (-) | grassland (+) |
| PC3 | grassland (+) | built up fraction (-), impervious density (-) | building height (-), traffic routes (-) |
| PC4 | shrubland (+) | diversity (+) | |
| PC5 | traffic routes (+) | diversity (+) | building height (-), built up fraction (-) |
| PC6 | traffic routes (+) | diversity (+), built up fraction (-), impervious density (-) | traffic routes (+), building height (-) |

Table B.1: Correlations between Urban Form Local features and Principal Components

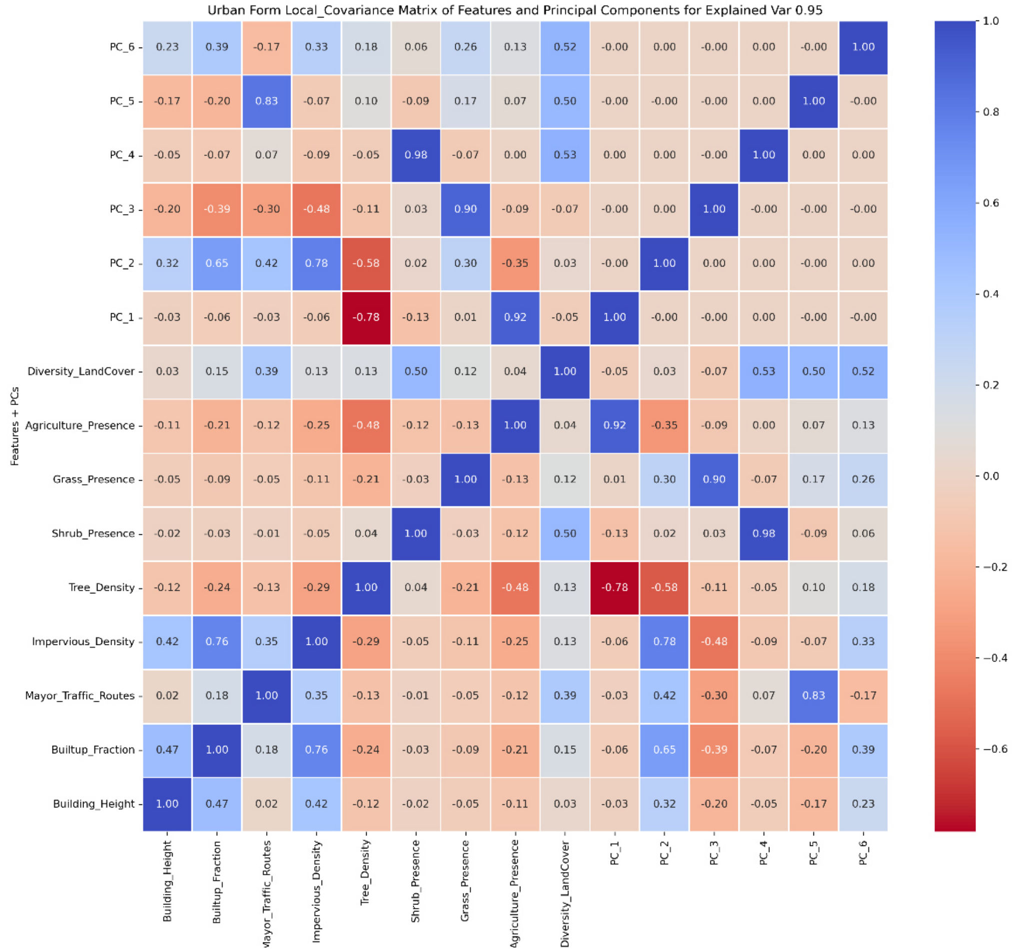


Fig. B.1: Urban Form Local PCA-features covariance matrix

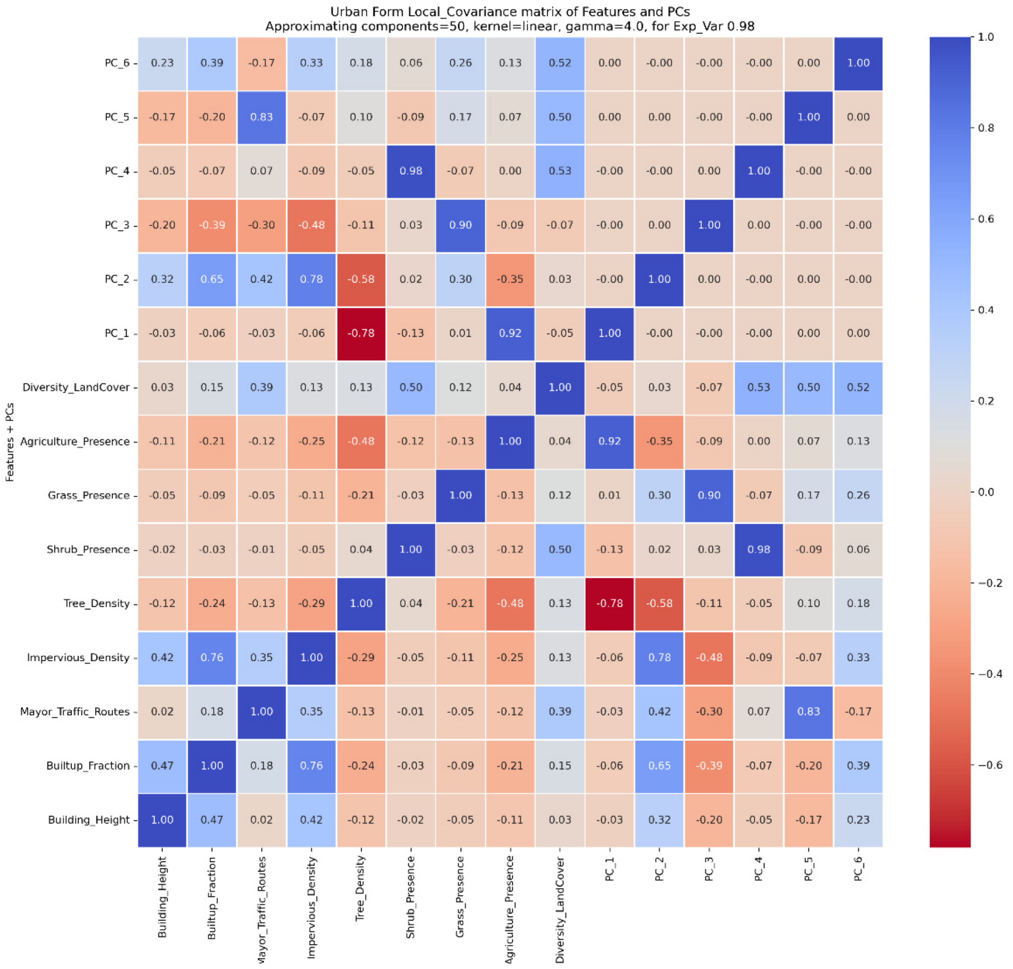


Fig. B.2: Urban Form Local KPCA-features covariance matrix

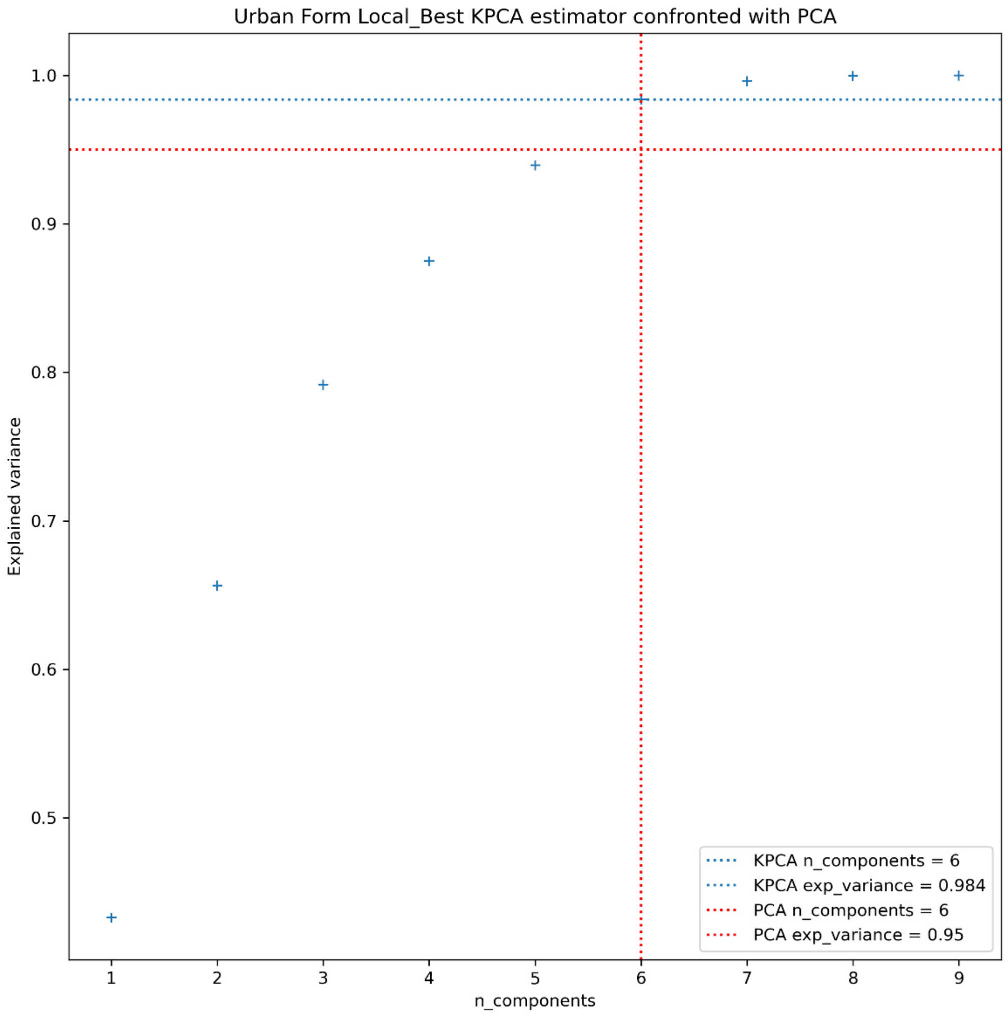


Fig. B.3: Urban Form Local best KPCA approximation confronted with PCA explained variance and number of components needed.

#### Urban Form Landscape

The analysis confronted the PCA and KPCA methods with similar results. The first explained 0.95 variance with 7 PCs (Fig. 1), while the best kernel of the latter explained 0.944 variance with 6 components (Fig. 2). Albeit with a very slim margin of difference, we went for the PCA set of components, as indicated from the confront plot (Fig. 3). We define the 7 PCs as indicated in Table B,2, ignoring contributions with covariance below 0.1, and referring to PC axis for the local scale. Shannon's Diversity Index is used to represent diversity in the window kernel, contrasting with local diversity. Contagion, instead, is used as a measure of aggregation within the kernel.

| Principal Component | Strong correlation (\|cov\| > 0.6) | Medium correlation (0.3 < \|cov\| <= 0.6) | Weak correlation (0.1 < \|cov\| <= 0.3) |
|---|---|---|---|
| PC1 | Diversity index (+), mean and std PC2 (+), std PC3 (+), std PC5 (+), std PC6 (+) | Mean PC1 (+), mean and std PC4 (+) | Mean PC3 (+), mean PC5 (-), mean PC6 (-) |
| PC2 | Mean PC1 (+) | Contagion (+), std PC1 (+) | Mean PC3 (+), mean PC4 (+) |
| PC3 | Contagion (-) | Mean PC1 (+), std PC1 (-), mean PC2 (+) | Std PC2 (-), Mean and std PC3 (-), mean and std PC4 (+) |

| | | | |
|---|---|---|---|
| PC4 | Mean PC3 (-) | Std PC1 (-), mean PC2 (+), std PC3 (-), std PC5 (+) | Contagion (+), mean PC6 (+) |
| PC5 | | Std PC3 (-), Mean and std PC4 (+) | Mean and std PC2 (-), Mean PC3 (-), mean and std PC5 (-), mean and std PC6 (-) |
| PC6 | | Std PC1 (-), mean and std PC3 (+) | Mean (+) and std PC2 (-), mean and std PC4 (+), std PC5 (-) |
| PC7 | Mean PC5 (-) | | Mean PC2 (+), Mean PC4 (-), std PC5 (-), mean (-) and std PC6 (+) |

Table B.2: Correlations between Urban Form Local features and Principal Components

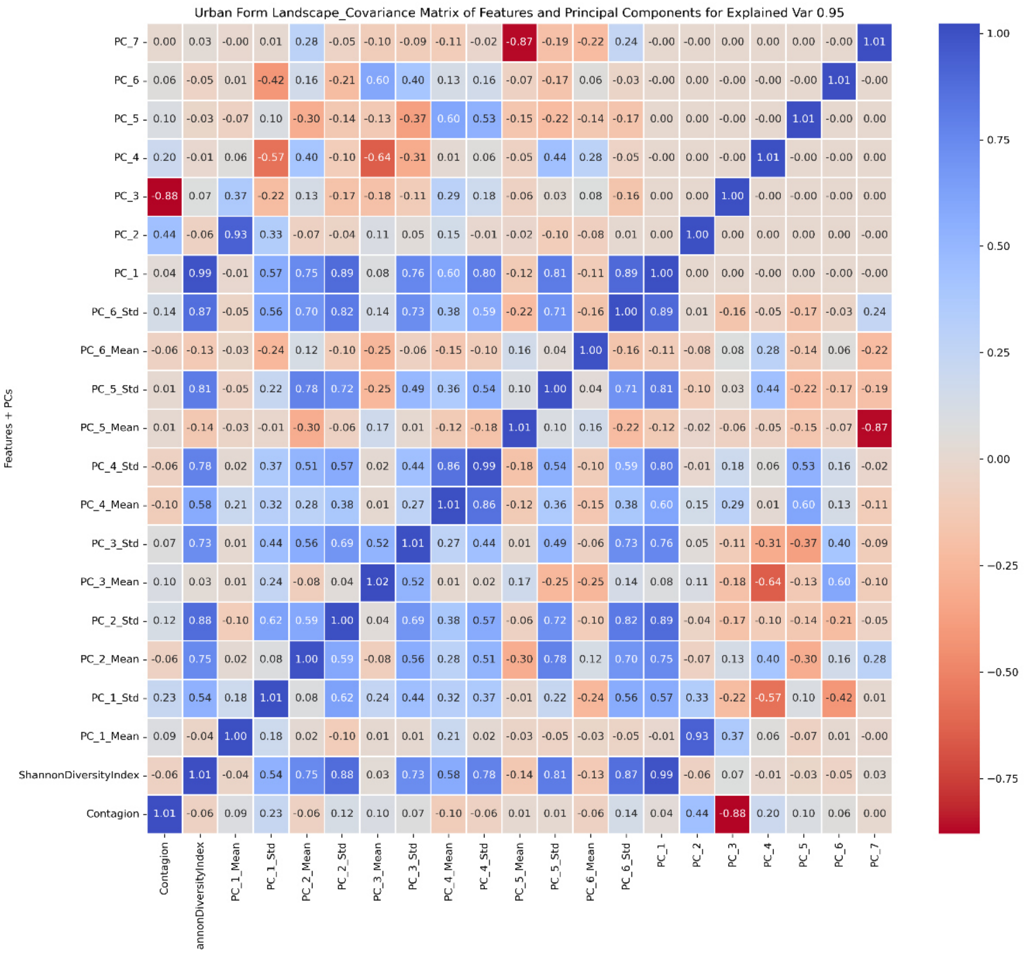


Fig. B.4: Urban Form Landscape PCA-features covariance matrix

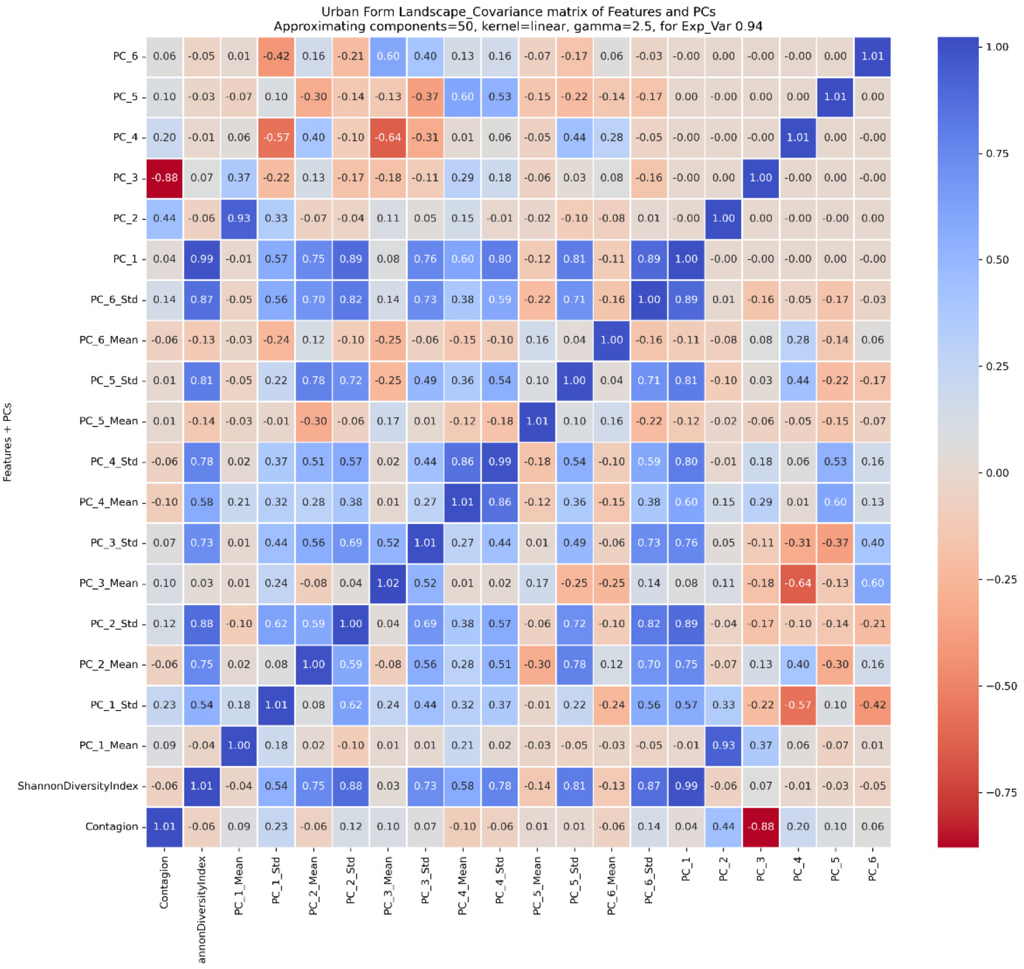


Fig. B.5: Urban Form Landscape KPCA-features covariance matrix

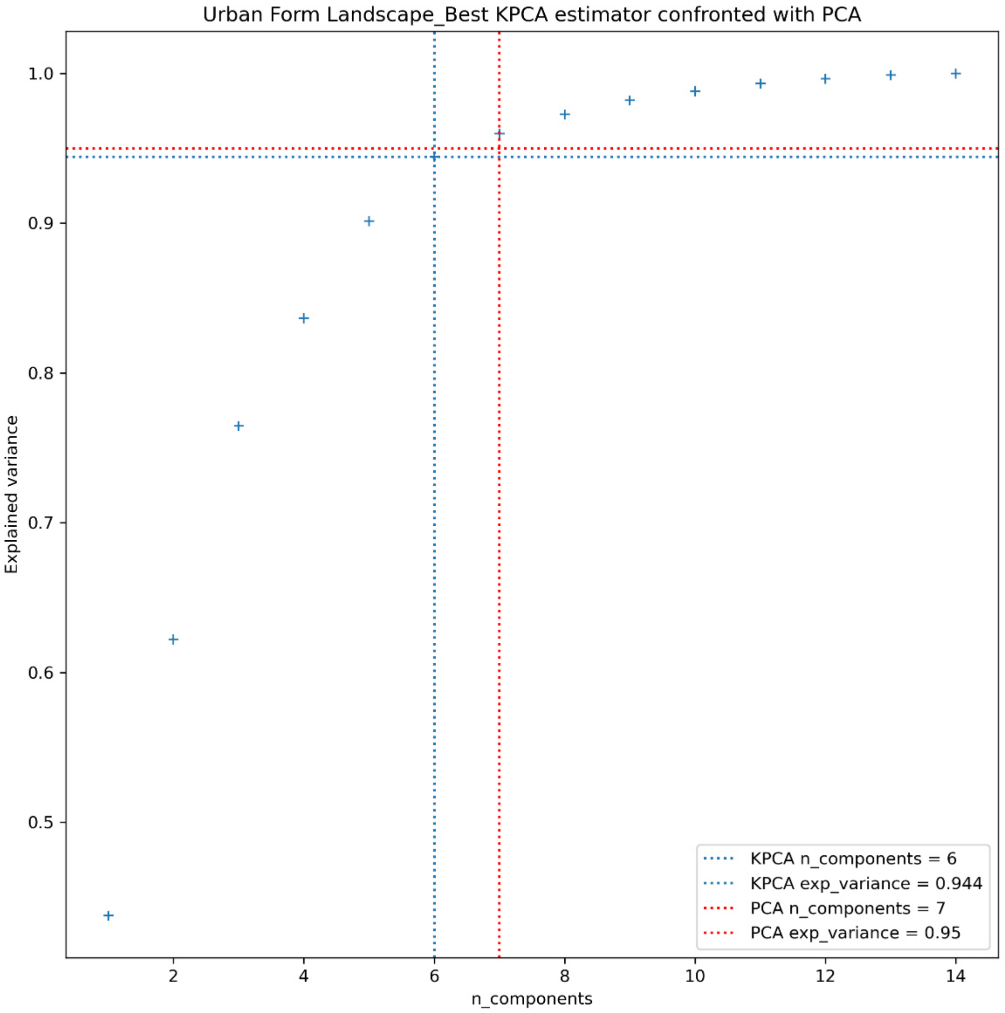


Fig. B.6: Urban Form Landscape best KPCA approximation confronted with PCA explained variance and number of components needed.

#### Anthropic Imprint

The analysis confronted the PCA and KPCA methods with similar results. The first explained 0.95 variance with 5 components (Fig. B.7), while the best kernel of the latter explained 0.969 variance with 6 components (Fig. B.8). Therefore, even though KPCA could explain more variation, we opted to use PCA since the minimum threshold of 0.95 variation could be explained with less components (Fig. B.9). We defined the 5 components in Table B.3, ignoring contributions with covariance below 0.1.

| Principal Component | Strong correlation (\|cov\| > 0.6) | Medium correlation (0.3 < \|cov\| <= 0.6) | Weak correlation (0.1 < \|cov\| <= 0.3) |
|---|---|---|---|
| PC1 | Population density (+), Age groups 0-15, 20-35, 40-60, 65+ (+) | Public green (+), preservation policy (-), public transport (-), nighttime light (+), heat emission (+) | Commercial area (+), industrial area (+), public service (+), |
| PC2 | Preservation policy (+) | | Age groups (+), recreation area (+) |
| PC3 | Recreation area (+) | Public green (+) | Heat emission (+) |
| PC4 | Public green (+) | | Population density (+), recreation area (-), cultural area (+), public |

| | | | |
|---|---|---|---|
| | | | transport (-), nighttime light (+), heat emission (+) |
| PC5 | Public transport (+) | | Public green (+) |

Table B.3: Correlations between Anthropic Imprint features and Principal Components

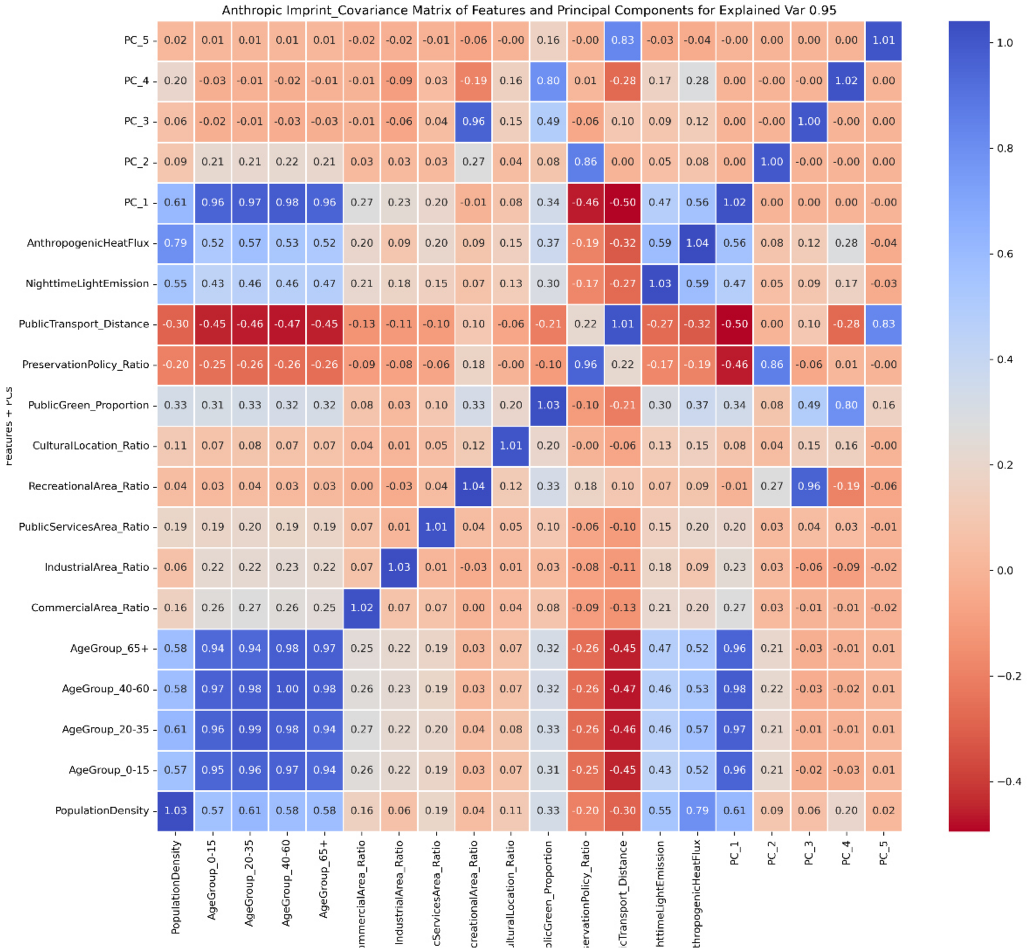


Fig. B.7: Anthropic Imprint PCA-features covariance matrix

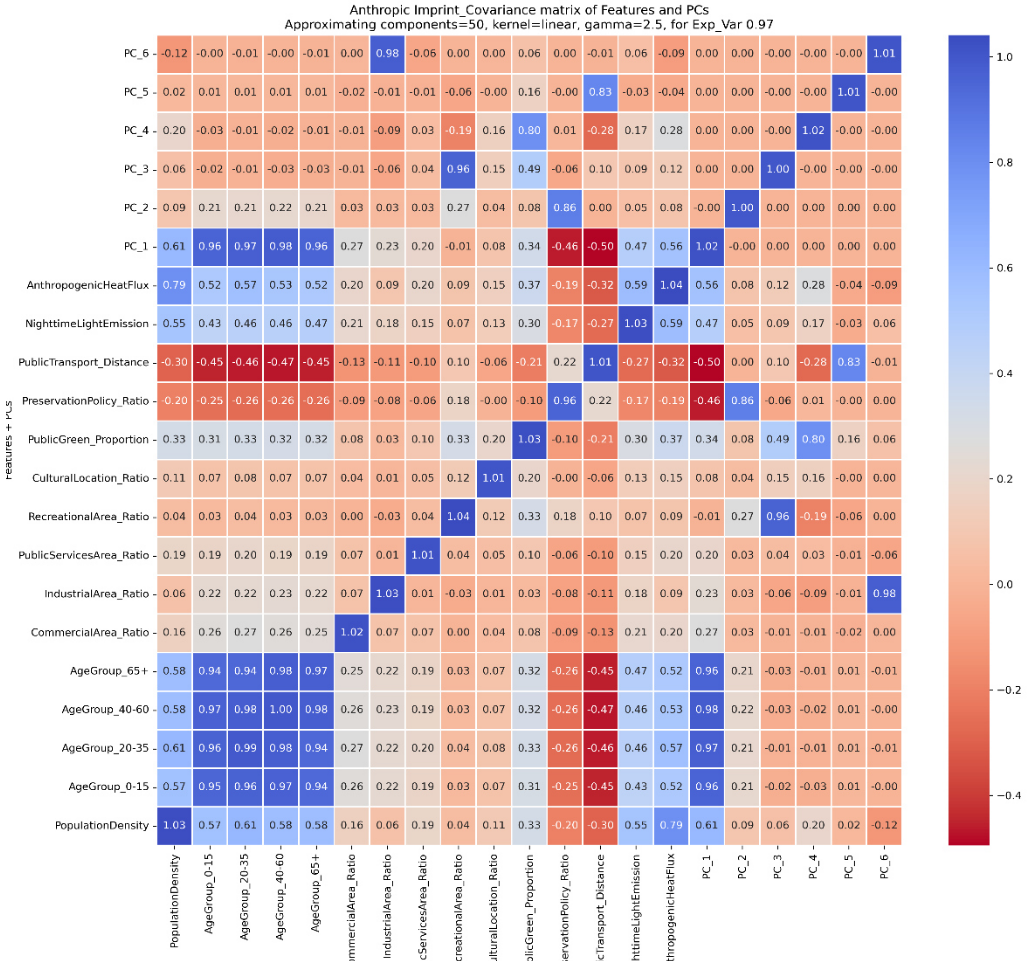


Fig. B.8: Anthropic Imprint KPCA-features covariance matrix

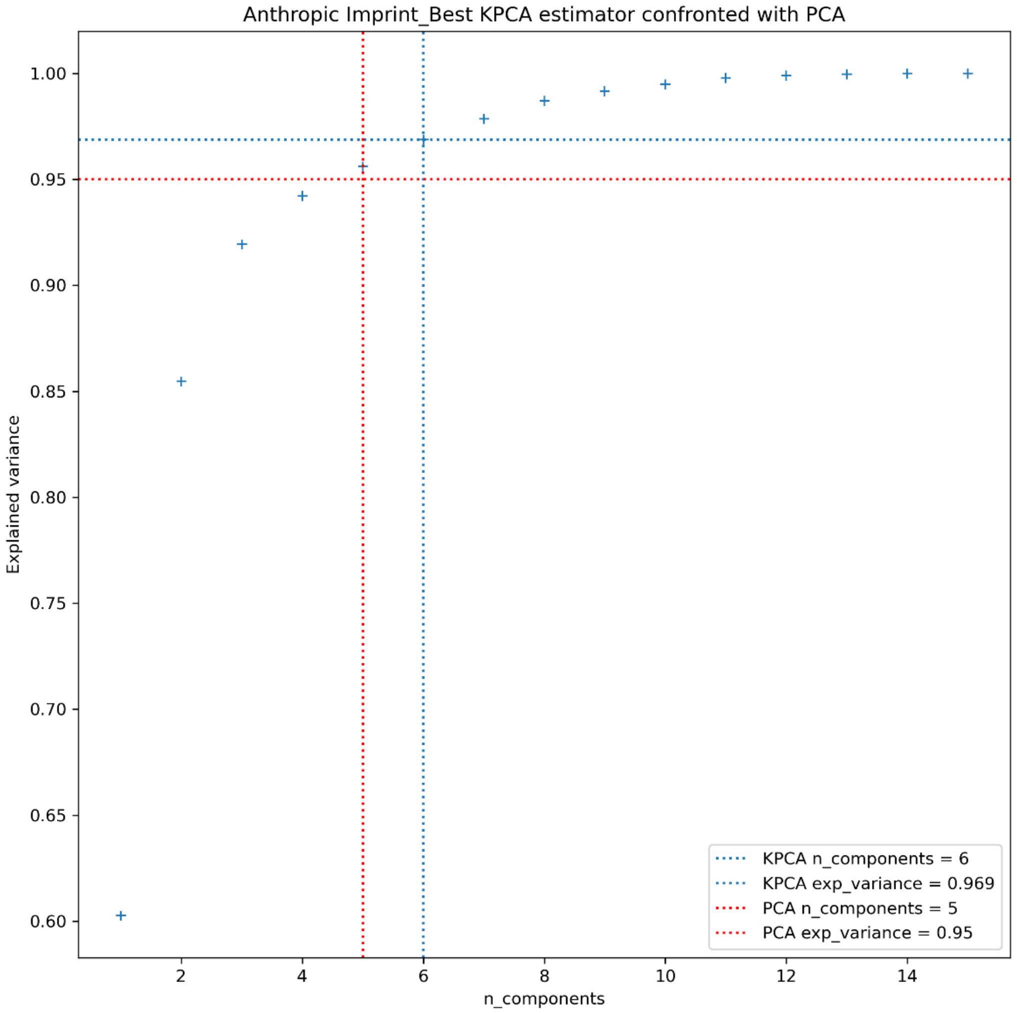


Fig. B.9: Anthropic Imprint best KPCA approximation confronted with PCA explained variance and number of components needed.

#### Biophysical Conditions

The analysis confronted the PCA and KPCA methods. The first explained 0.95 variance with 8 components (Fig. B.10), and the best kernel of the latter explained 0.908 variance 6 components (Fig. B.11). Therefore, having not met the minimum threshold, we discarded the KPCA components in favour of the PCA (Fig. B.12). We define the 8 PCs in Table B.4, ignoring contributions with covariance below 0.1. We noted that surface water and soil wetness behaved with perfect correspondence, similarly to wind speed at 10m and 200m.  Values above 1 should be considered as 1 and comes from approximations of the algorithm.

| Principal Component | Strong correlation (\|cov\| > 0.6) | Medium correlation (0.3 < \|cov\| <= 0.6) | Weak correlation (0.1 < \|cov\| <= 0.3) |
|---|---|---|---|
| PC1 | Altitude mean (+), temperature max and min relative and absolute (-), precipitations absolute (+) | Precipitations relative (+) | Altitude std (+), slope (+), aspect mean (+), curvature std (+), wind speed (-) |
| PC2 | Altitude std (+), slope (+), curvature std (+) | Precipitations absolute and relative (+) | Soil wetness (-), temperature max and min relative (-) max and min absolute (+) |

| | | | |
|---|---|---|---|
| PC3 | | Aspect (-), temperature max and min relative (-), precipitations absolute (-) and relative (+),wind speed (+) | Slope mean (-), curvature std (-) |
| PC4 | Aspect std (-) | Permanent and temporary water (+), aspect mean (-), temperature max and min relative (+) | Precipitations absolute (+), wind speed (-) |
| PC5 | Permanent and temporary water (+) | | Aspect (+), wind speed (+) |
| PC6 | Aspect mean (+) | Aspect std (-) | |
| PC7 | Soil wetness (+) | Wind speed (+) | Precipitations absolute (+) |
| PC8 | | Soil wetness (+), precipitations relative (-), wind speed (-) | Precipitations absolute (-) |

Table B.4: Correlations between Anthropic Imprint features and Principal Components

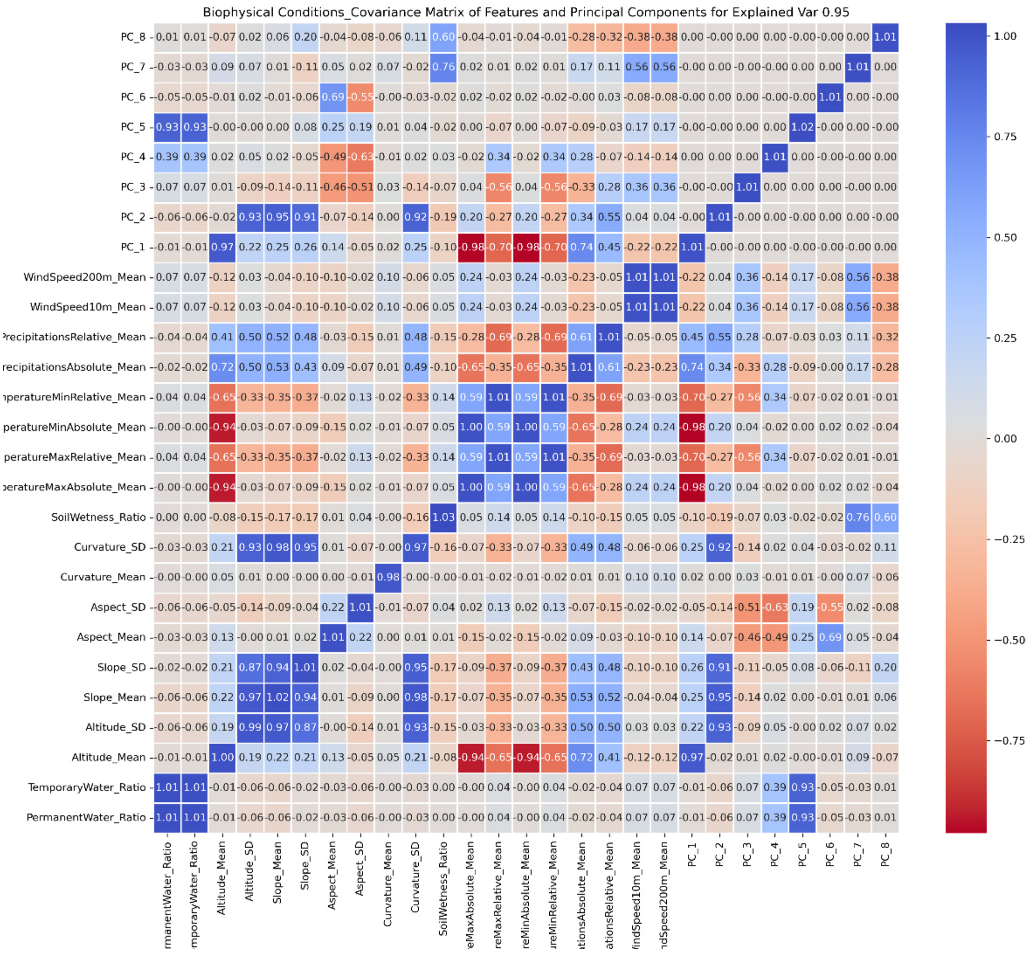


Fig. B.10: Biophysical Conditions PCA-features covariance matrix

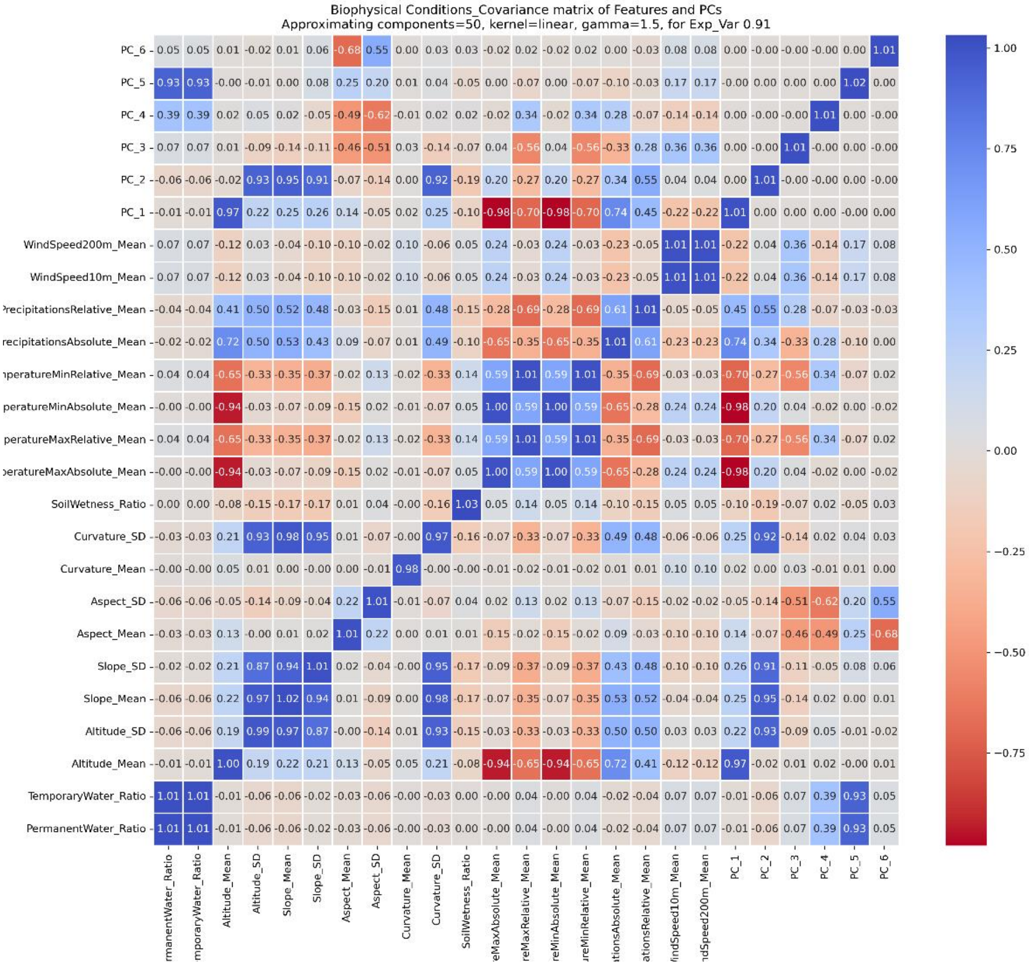


Fig. B.11: Biophysical Conditions KPCA-features covariance matrix

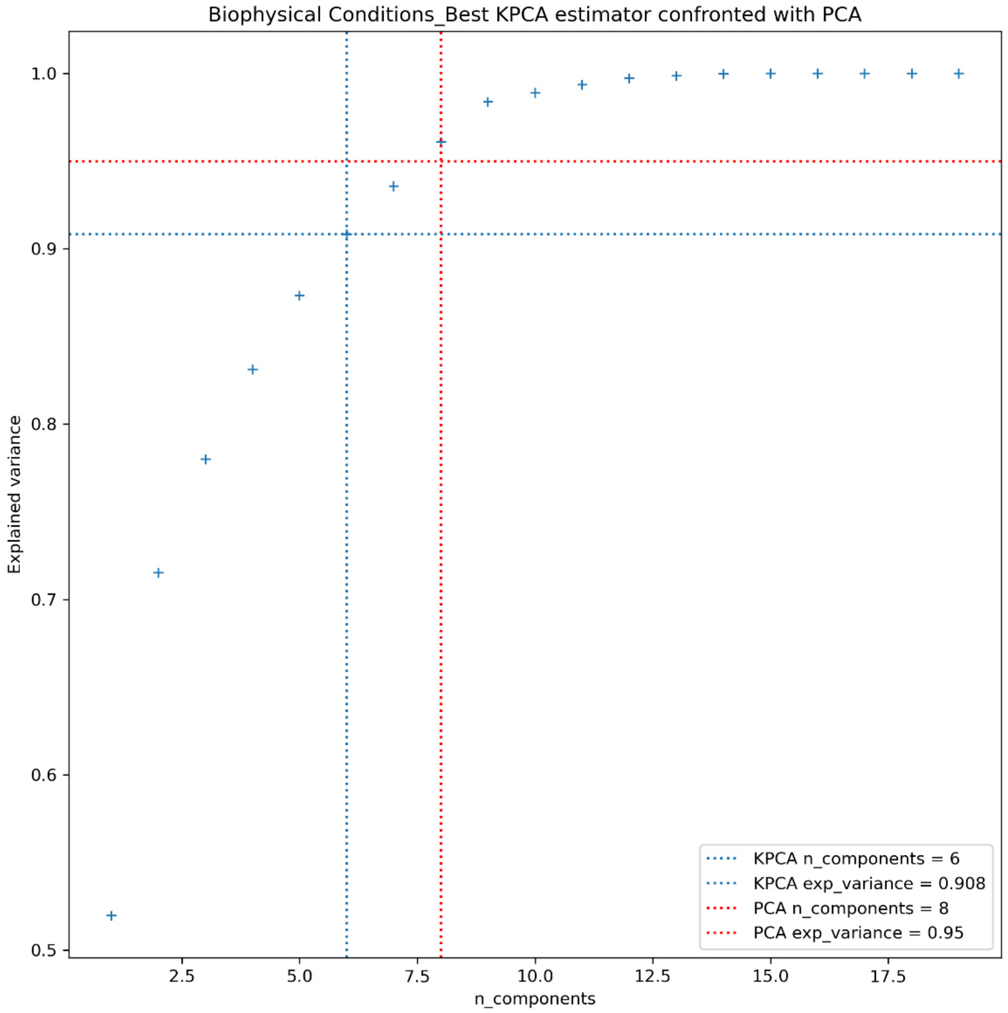


Fig. B.12: Biophysical Conditions best KPCA approximation confronted with PCA explained variance and number of components needed.
