## Supplementary material for "Urban Cohabscapes: exploring European Co-Habitative landScapes diversity, in the ECOLOPES framework": Annex C

### Annex C. Unsupervised Classification scores

We initially tested the Hierarchical Clustering approach, as agglomerative clustering showed predominantly among classifications of urban landscapes (Paumelle et al., 2023), as well as DBSCAN, HBDSCAN, OPTICS for their increased flexibility and reduced time complexity, and Random Forest and Unsupervised Neural Networks for their expected increase in clustering quality. We noted that the computation complexity was still a problem, infact our sub-classification datasets were defined by over 60’000’000 cells given the extensive spatial boundaries and resolution. We finally tested the KMeans family of algorithms and confronted different flavours. we discarded the Fuzzy and Bisecting KMeans as they showed clear unfavourable performance against the traditional Lloyd implementation. We then limited to the Lloyd, Elkan and MiniBatch, using sklearn implementations for all three.

The sub-classification are processed through three different flavors of KMeans: Lloyd, Elkan, MiniBatch, using sklearn implementations. To identify the strongest classification, and selecte the number of distinct classes (k), we used three different scores suggested by literature: Silhouette score (SIL), Calinski-Harabasz (CH) and Davies-Bouldin (DB) (Palacio-Niño and Berzal, 2019). Our goal is to maximise clustering performance with an automatic criterion for k selection. For each possible k (between 6 and 30) we calculate each score; then, we take the overall maximum score and calculate the difference between the k-th score and the maximum score. We normalize and sum the three differences together and locate the best solution as the nearest between the three different maximums. We do this for each k, for each of the three algorithms, and we plot the results identifying the maximum value for the Silhouette and Davies-Bouldin scores, and the minimum for the Calinski-Harabasz. For validating the prediction, we plot the inertia and use the Elbow Method to visualize the curvature of inertia for the growing number of k.

#### Urban Form Local

We run through the classifier two times, extracting a portion of data referred to the built environment to process additional classes. In the first run (Fig. C.1, C.2, C.3) we identify the use of Elkan with k=10 and scores of SIL=0.819, CH=1.742e7, DB=0.485, while in the second run (Fig. C.4, C.5, C.6) we select Elkan with k=4 and SIL=0.689, CH=1.631e6, DB=0.619.

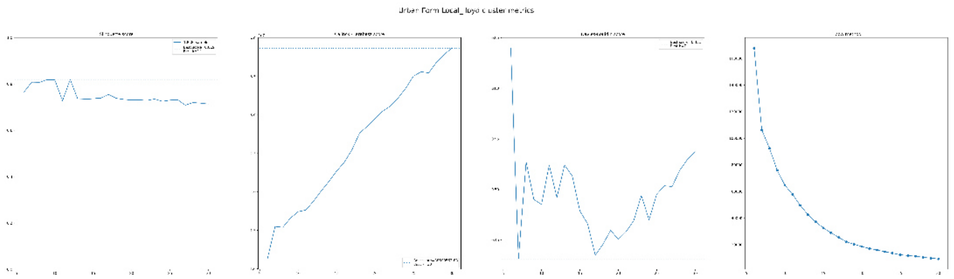

Fig. C.1: Urban Form Local (first run) Lloyd KMeans score plots.

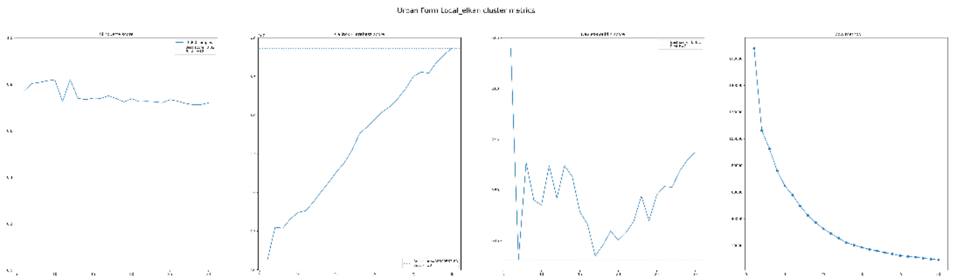

Fig. C.2: Urban Form Local (first run) Elkan KMeans score plots.

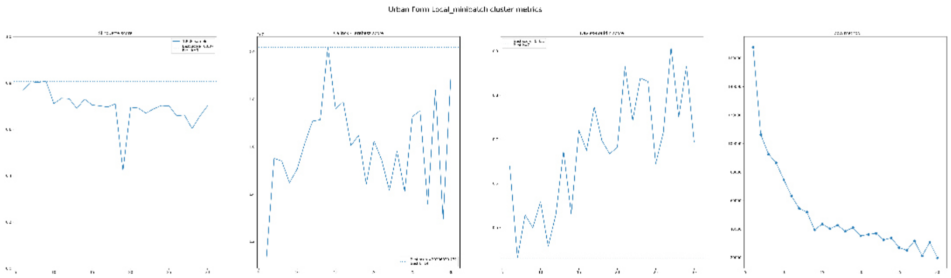


Fig. C.3: Urban Form Local (first run) MiniBatch KMeans score plots.

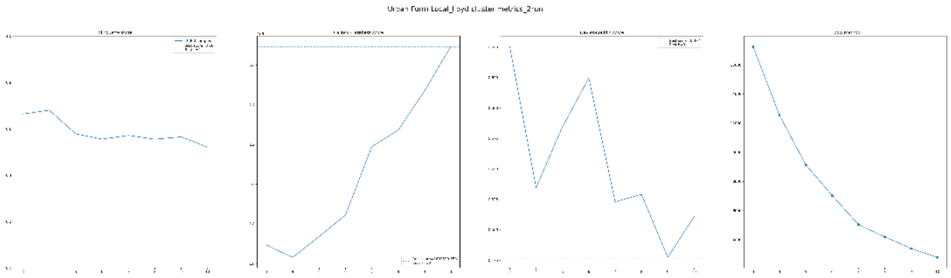


Fig. C.4: Urban Form Local (second run) Lloyd KMeans score plots.

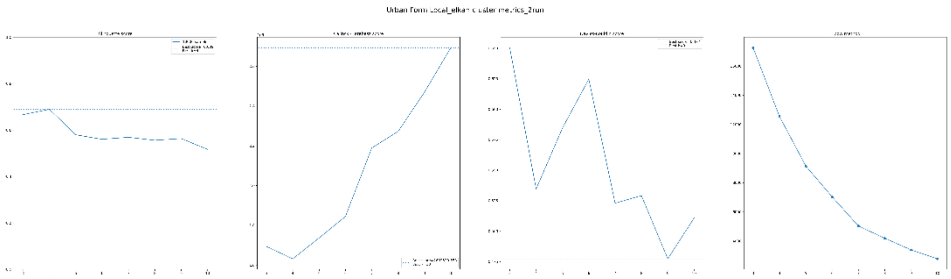


Fig. C.5: Urban Form Local (second run) Elkan KMeans score plots.

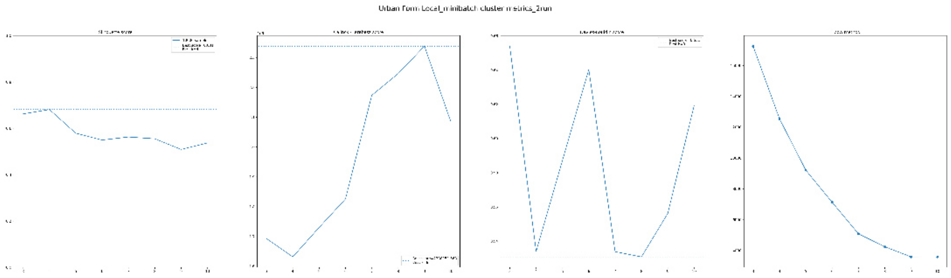


Fig. C.6: Urban Form Local (second run) MiniBatch KMeans score plots.

#### Urban Form Landscape

We run through the classifier one time and select the best algorithm and k value in Elkan with k=6, with scores SIL=0.440, CH=3.035e6, DB=0.990. (Fig. C.7, C.8, C.9).

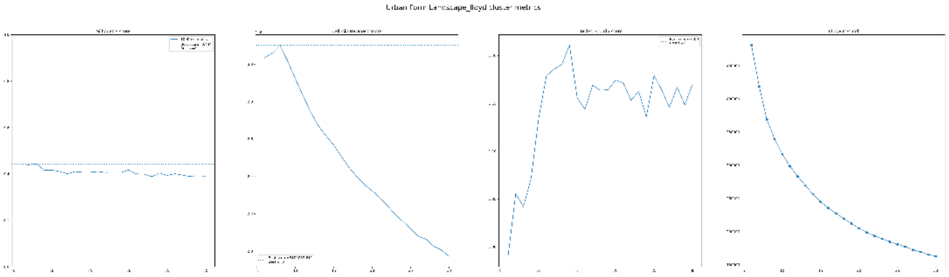

Fig. C.7: Urban Form Landscape Lloyd KMeans scores plots.

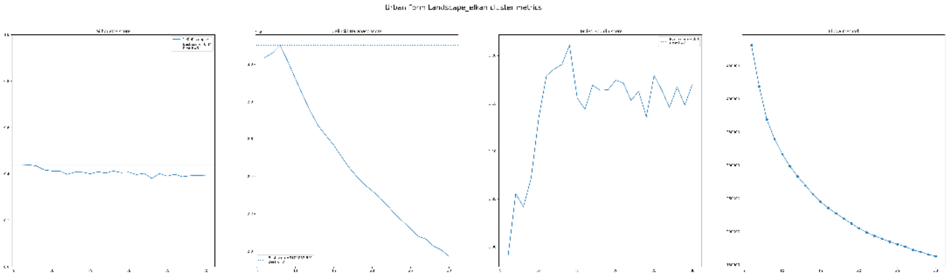

Fig. C.8: Urban Form Landscape Elkan KMeans scores plots.

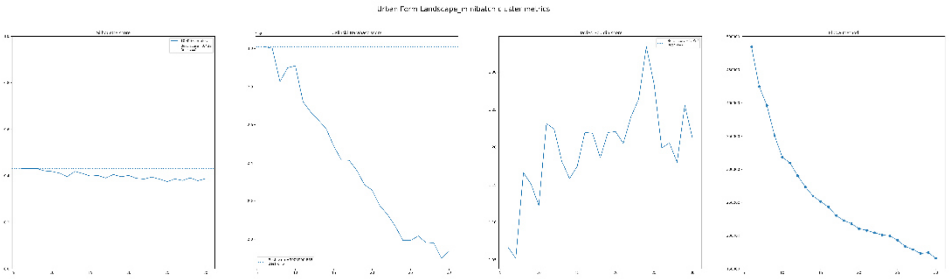

Fig. C.9: Urban Form Landscape MiniBatch KMeans scores plots.

#### Anthropic Imprint

We classify one time and from the gridsearch we select the best combination of algorithm and k value in Elkan with k=6 and SIL=0.662, CH=6.846e6, DB=0.723 (Fig. C.10, C.11, C.12).

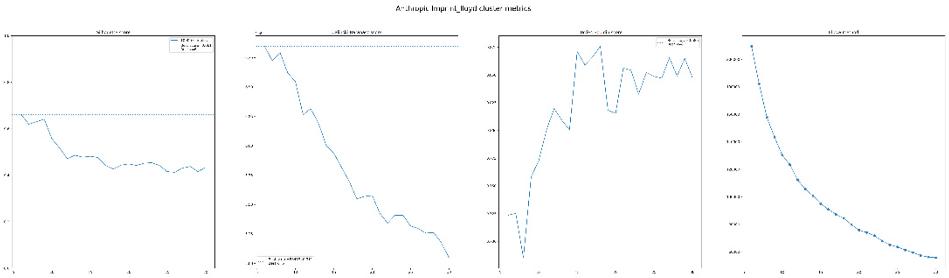

Fig. C.10: Anthropic Imprint Lloyd KMeans scores plots.

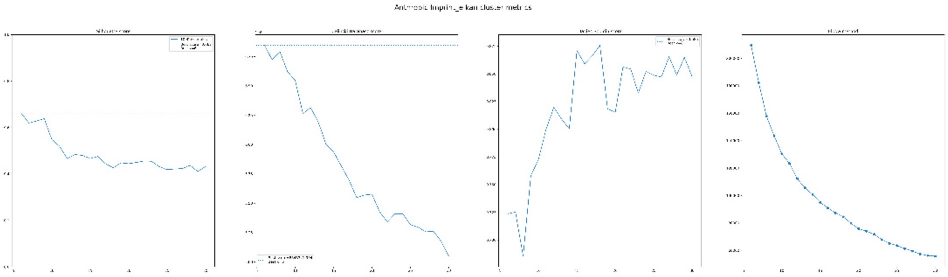

Fig. C.11: Anthropic Imprint Elkan KMeans scores plots.

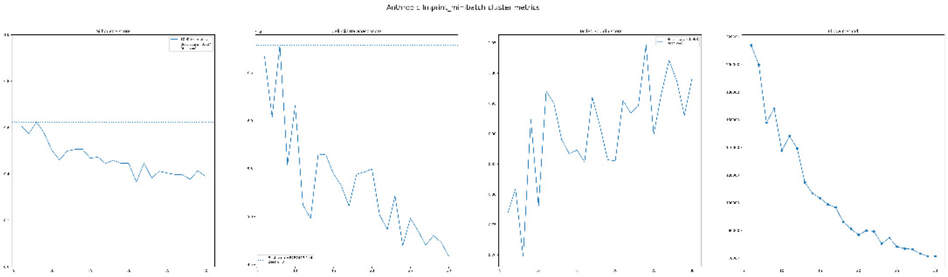

Fig. C.12: Anthropic Imprint MiniBatch KMeans scores plots.

#### Biophysical Conditions

We run the classifier pipeline one time, and select the combination of Lloyd with k=6 and SIL=0.371, CH=1.713e6, and DB=1.318 (Fig. C.13, C.14, C.15).

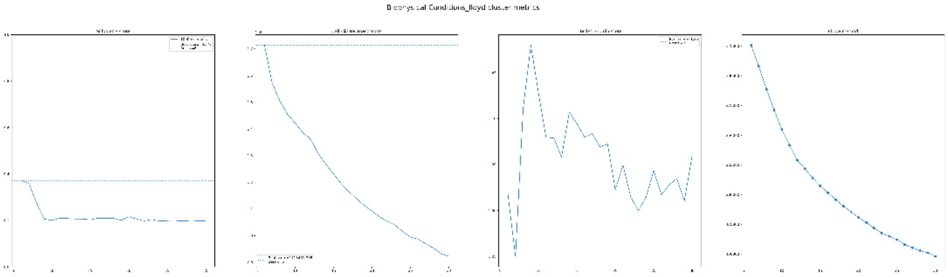

Fig. C.13: Biophysical Conditions Lloyd KMeans scores plots.

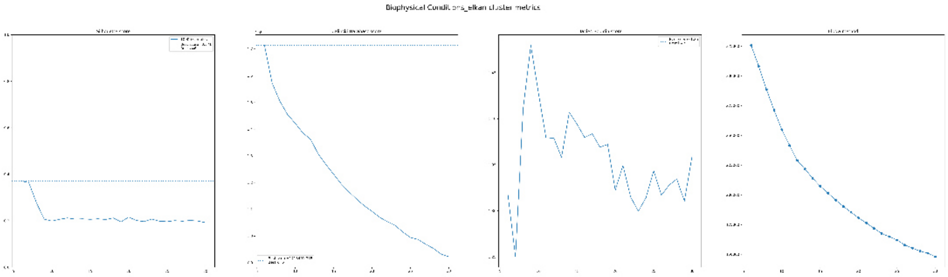

Fig. C.14: Biophysical Conditions Elkan KMeans scores plots.

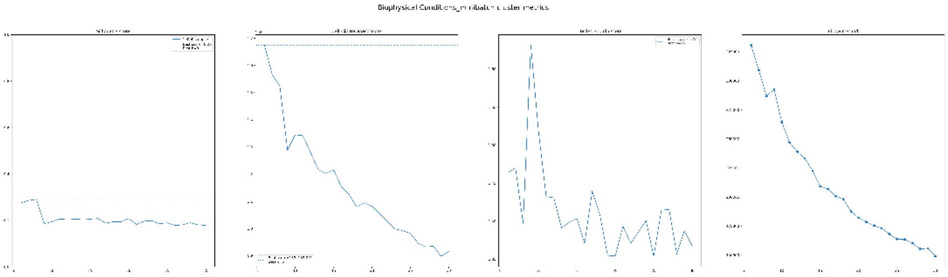


Fig. C.15: Biophysical Conditions MiniBatch KMeans scores plots.
